# Structured and Target-Specific Development of Cortico-Cortical Connectivity in the Mouse Visual Cortex

**DOI:** 10.1101/2025.11.03.686379

**Authors:** Matthew W. Jacobs, John M. Ratliff, Alec L.R. Soronow, Jordan A. Nichols, Hylen T. James, Jorin A.G. Eddy, Adam M. Murray, Euiseok J. Kim

## Abstract

The mammalian cortex exhibits highly stereotyped long-range connectivity, yet the developmental principles that specify precise cortico-cortical projection patterns remain poorly defined. Two dominant models propose that target specificity arises either from early inter-regional exuberant outgrowth followed by pruning, or through initially directed axonal targeting. To resolve this, we systematically mapped the postnatal development of V1 cortico-cortical projection neurons (CCPNs) to eleven higher visual areas (HVAs) in mice using rapid and complementary retrograde, anterograde, and single-cell tracing methods. We found that V1→HVA connectivity develops via spatiotemporally staggered axon extension and pruning programs, aligned with target position along the medial-lateral axis. Reciprocal HVA→V1 feedback emerges concurrently and is refined over time, yielding gradually aligned bidirectional connectivity. Notably, both multiplexed retrograde tracing and MAPseq-based single-cell profiling revealed that individual V1 neurons initialize and retain specific projection motifs with limited variation over development, arguing against global exuberance followed by selective, inter-areal pruning. Instead, our findings support a directed guidance model, in which distinct V1 CCPN subtypes establish selective projection patterns early, followed by local, target-dependent refinement. This structured yet heterogeneous developmental strategy provides an anatomical framework for how precise long-range cortical networks emerge.

**HIGHLIGHTS:**

- V1→HVA connections form via directed axonal targeting, establishing motifs early with little variation
- Medial targets are innervated earlier and refine gradually, lateral targets later and rapidly
- Feedforward and feedback V1-HVA circuits emerge concurrently
- Bidirectional like-to-like V1–HVA connectivity refines across development

## INTRODUCTION

In the mammalian cerebral cortex, the transformation of sensory inputs into coherent percepts is mediated by inter-areal connectivity between hierarchically organized cortical regions.^1–3^ The proper development of cortico-cortical connections is essential for neocortical functions such as hierarchical computation, multisensory integration, and the control of complex behavior. Disruptions to this process are implicated in neurodevelopmental and psychiatric disorders characterized by aberrant inter-areal connectivity, including autism spectrum disorder and schizophrenia.^4,5^ Understanding how these circuits form at cellular resolution is critical to explaining how long-range cortical networks acquire their mature functional architecture.

In mice, cortico-cortical projection neurons (CCPNs) in the primary visual cortex (V1) project to at least ten anatomically and functionally distinct higher visual areas (HVAs), forming precise circuits that support visual information processing.^6–9^ In the adult brain, CCPNs show clear target specificity: some project to a single HVA as dedicated information channels, while others innervate multiple HVAs in a non-random, structured manner, functioning as broadcasting channels.^10^ Furthermore, reciprocal feedback from HVAs often targets the same subpopulations of V1 neurons that project to them, forming “like-to-like” connectivity motifs.^7,11^ However, the developmental basis of this specificity remains poorly understood.

Two main developmental models have been proposed to explain how CCPNs acquire long-range target specificity.^12,13^ The exuberant growth and pruning model suggests that immature neurons initially send projections to multiple targets, followed by activity-dependent pruning of incorrect or redundant axonal branches, resulting in refined adult patterns.^14–18^ In contrast, the directed guidance model posits that target specificity is largely established early, driven by intrinsic genetic programs and molecular cues that guide target-directed axonal growth.^19–21^ Distinguishing between these models in the context of multi-area cortico-cortical circuit development has been challenging, partly due to technical limitations in tracing the full axonal output of individual neurons across time with sufficient spatial and temporal resolution.

To address this, we systematically characterized the postnatal development of ipsilateral cortico-cortical connectivity between V1 and HVAs using a combination of chemical and viral, anterograde and retrograde, anatomical tracing strategies, including cholera toxin subunit B (CTB, retrograde), vesicular stomatitis virus (VSV, anterograde), AAV-retrograde vectors, and barcoded Sindbis virus (anterograde). Through rapid dual and sequential retrograde labeling, rapid bulk anterograde tracing, and quantitative histology, we found that V1 neurons develop their axonal projections to HVAs in a spatiotemporally heterogeneous manner, depending on target position along the medial-lateral axis. Retrograde tracing revealed that CCPNs projecting to medial and lateral HVAs emerge in a biased temporal sequence, with medial targets innervated days earlier than lateral ones. Bulk anterograde tracing confirmed that medial circuits mature gradually, while lateral circuits exhibit transient overgrowth followed by rapid refinement. Our extended survival experiments ruled out large-scale cell death, indicating that axon pruning in the designated target area is a developmental mechanism driving circuit refinement. Additionally, feedback projections from HVAs to V1 emerged with less spatial and temporal heterogeneity than feedforward projections from V1 to HVAs, resulting in the progressive development of reciprocal ‘like-to-like’ connectivity through the gradual alignment of feedforward and feedback pathways. Finally, we used barcoded Sindbis virus with the MAPseq (Multiplexed Analysis of Projections by Sequencing) platform^22^ to label and quantify the projection motifs of thousands of individual V1 CCPNs across developmental timepoints. MAPseq revealed that the number of target areas innervated by each neuron, the complexity of their projection patterns, and the frequency of specific motifs remained largely consistent across development.

Together, our findings support a directed guidance model in which target specificity, temporally staggered axon innervation, and pruning within defined target areas contribute to circuit formation. Rather than widespread inter-regional axonal reorganization, the development of inter-areal cortical connectivity is shaped by the selective emergence of projection motifs.

## RESULTS

### Axonal targeting from V1 to AL and PM occurs via directed projection during development

V1 cortico-cortical projection neurons (CCPNs) communicate with higher visual areas (HVAs) using at least two distinct modes: dedicated projections to a single target area, or broadcasting projections to multiple areas following specific patterns^7,10^ (Figure 1A). As an example of the dedicated mode, two largely non-overlapping V1 CCPN populations project to either AL or PM (V1→AL or V1→PM CCPNs, respectively), without sharing axon collaterals across these targets, and are known to exhibit distinct visual response properties.^7,8,23^

**Figure 1.**
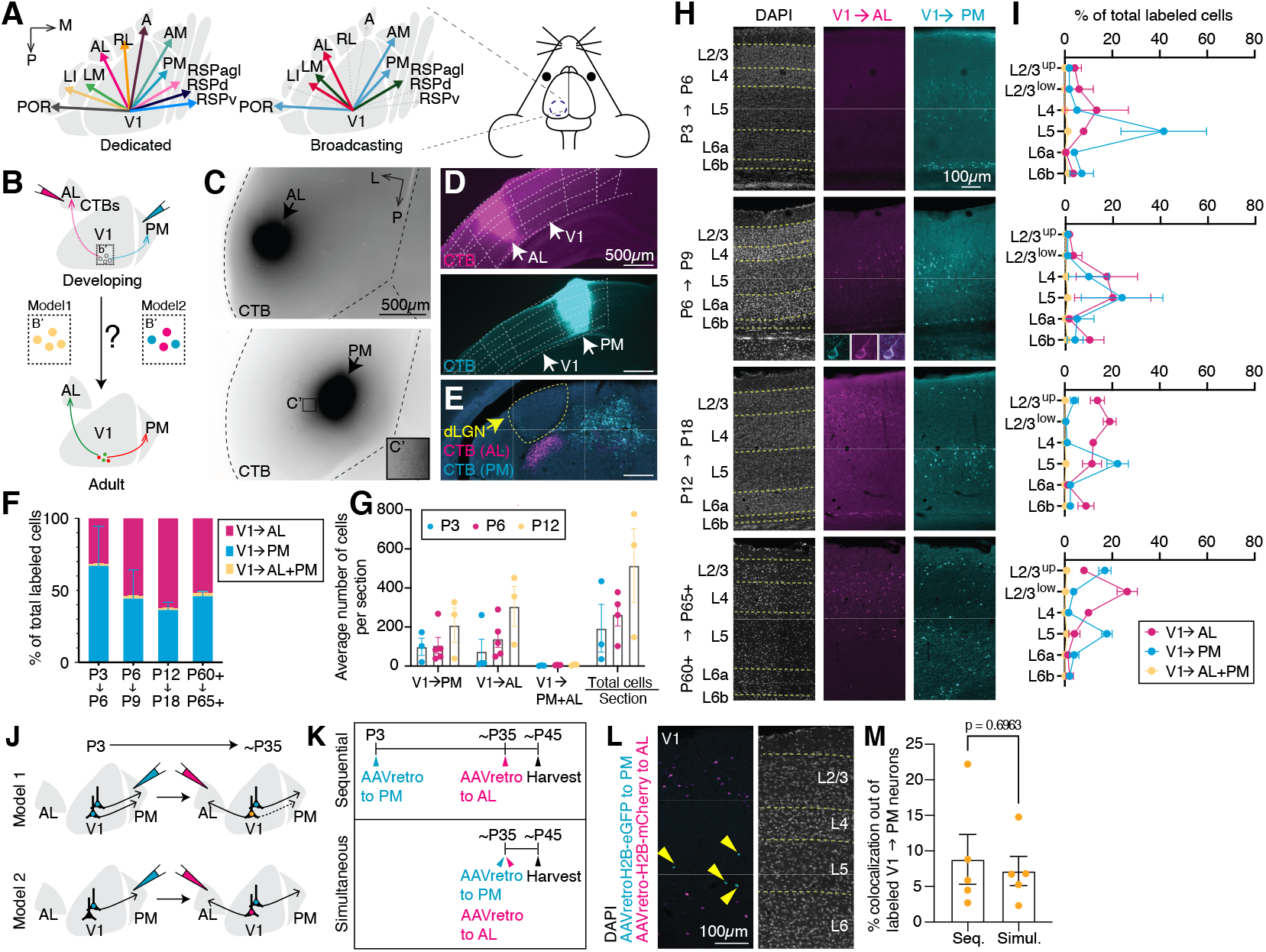
Directed axonal projection from V1 to higher visual areas (HVAs), AL and PM, during development. **(A)** Schematic illustrating two modes of long-range axonal projection from V1 to HVAs in the adult brain. In the dedicated mode, cortico-cortical projection neurons (CCPNs; arrows) project to a single HVA target area. In the broadcasting mode, CCPNs (arrows) extend axons to multiple HVAs in distinct patterns. **(B)** Strategy for projection-based neuronal labeling using fluorophore-conjugated cholera toxin subunit B (CTB) to differentiate between two V1→AL and V1→PM projection models across developmental time points. **(C)** Dorsal view of P6 brains showing precise CTB injections into AL (top) and PM (bottom). **(C’)** Magnified view of the inset location for CTB-labeled V1→PM CCPNs (black). **(D)** Representative confocal images of coronal sections from a P6 brain showing selective CTB labeling in AL (magenta) and PM (cyan) without spillover into V1. **(E)** A confocal image showing the absence of CTB-labeled neurons in the dorsal lateral geniculate nucleus (dLGN; yellow) and the presence in the lateral posterior nucleus (LP), confirming selective AL and PM targeting at P3. **(F)** Quantification of singly labeled V1→AL (magenta), V1→PM (cyan), and dual-labeled V1→AL+PM (yellow) CCPNs across developmental stages. **(G)** Average number of CTB-labeled CCPNs per section over developmental time. **(H-I)** Confocal images and laminar distributions of CTB-labeled V1→PM (cyan) and V1→AL (magenta) CCPNs at different developmental stages. **(J)** Schematic of two models for temporally distinct V1→AL and V1→PM axonal projection development. **(K)** Viral strategy using AAVretro to distinguish between these models in **J. (L)** Confocal images showing AAVretro-labeled nuclei of V1→AL and V1→PM CCPNs in V1 at P45. **(M)** Quantification of the proportions of double-labeled neurons among V1→PM-labeled neurons in the sequential (Seq.) and simultaneous (Simul.) injection groups. Student’s t tests. For CTB experiments, n = 3 mice for P3→P6, n = 4 for P6→P9, n = 3 for P12→P18, and n = 3 for P60+→P65+. For AAVretro experiments, n = 5 mice for P3→P35→P45 Exp. group and n = 5 for P35→P45 control group. Data are represented as mean ± SEM.

To investigate how this target-specific projection pattern arises during development, we injected Alexa Fluor 555 (magenta) or 647 (cyan)-conjugated cholera toxin subunit B (CTB) into AL and PM, respectively, at postnatal days P3, P6, P12, and P60+ (Figure 2B). Then, we quantified the numbers and proportions of V1→AL and V1→PM CCPNs 3–6 days after injection (referred to hereafter as P3→P6, P6→P9, P12→P18, and P60+→P65+), including the degree of dual labeling observed in each group (Figures 1A-B). If V1 CCPNs initially extend axons exuberantly to multiple areas, a higher proportion of neurons co-labeled for both AL and PM projections would be expected during early developmental stages compared to adulthood (Model 1, Figure 1B’). In contrast, if CCPNs form specific projections early on, the proportion of dual-labeled cells would remain low (Model 2, Figure 1B’).

**Figure 2.**
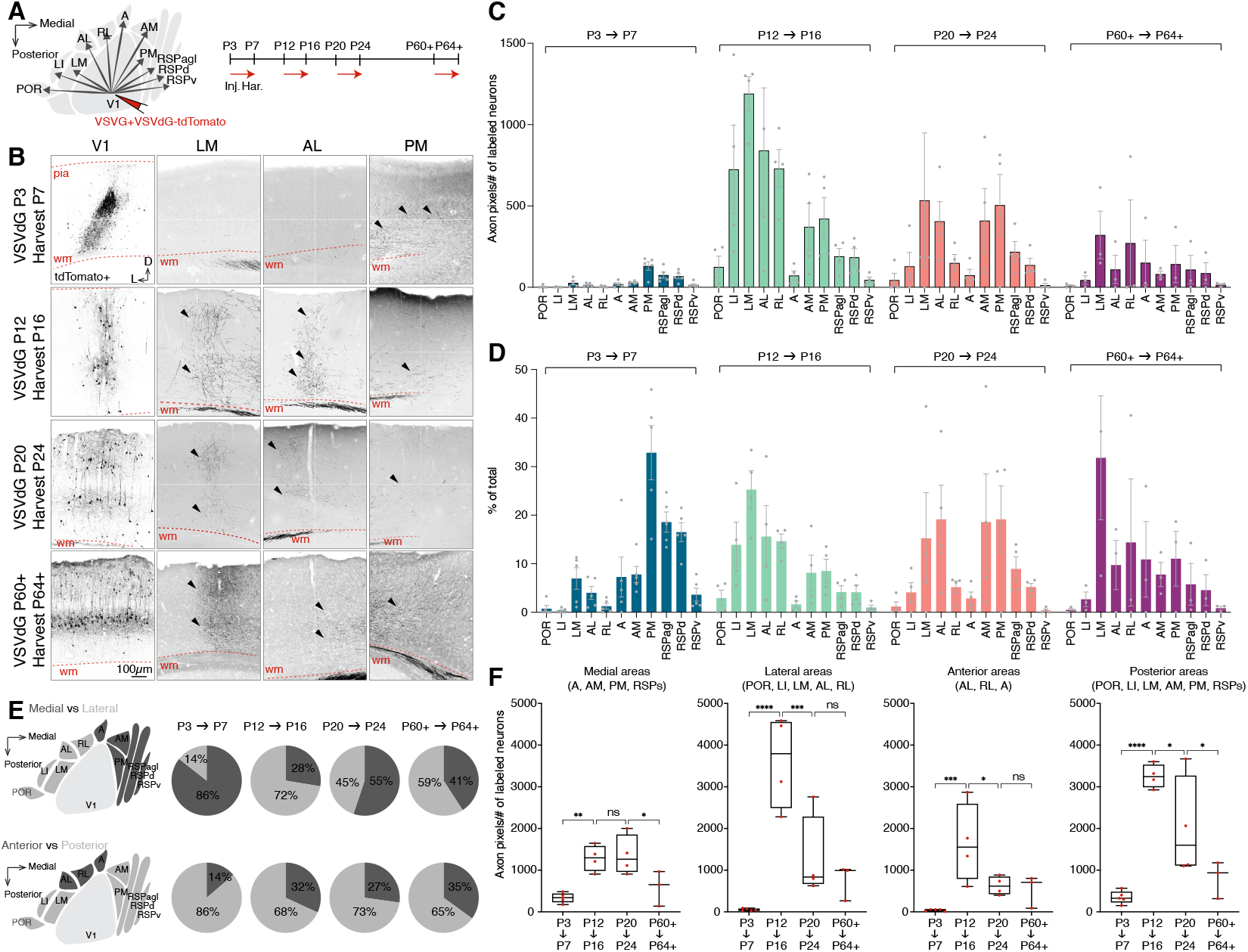
Reconstruction of long-range axonal projections from V1 to eleven cortical areas across postnatal stages. **(A)** Schematic of the experimental strategy. VSVG+VSVdG-tdTomato was injected into the center of V1 at various developmental and adult time points. Brains were collected 4 days post-injection for analysis. **(B)** Representative confocal images of coronal brain sections showing V1 injection sites and tdTomato+ neurons (leftmost column) and tdTomato+ long-range axons (arrows) projecting to multiple HVAs: LM, AL, and PM. **(C-D)** Quantification of axonal labeling in eleven cortical areas across development. **(C)** tdTomato+ axon pixel counts normalized per labeled V1 CCPN, and **(D)** the percentage of total tdTomato+ axonal output represented in each area. **(E)** Pie charts illustrating the distribution of tdTomato+ axons in medial versus lateral HVAs (top) and anterior versus posterior HVAs (bottom) at different developmental and adult stages. **(F)** Comparative analyses of tdTomato+ axon pixels per labeled V1 CCPN across developmental stages in medial, lateral, anterior, and posterior target groups. n = 5 mice for P3→P7, n = 4 for P12→P16, n = 4 for P20→P24, and n = 3 for P60+→P64+. one-way ANOVA with uncorrected Fisher’s LSD; *p < 0.05, **p < 0.01, ***p < 0.001, ****p < 0.0001. Data are shown as mean ± SEM.

To ensure the specific targeting of the higher visual area at early postnatal stages, we screened brains by examining whole-mount and coronal sections of the cortex and thalamus (Figures 1C-D). Injections into AL and PM were considered accurate only if there was no spillover into adjacent regions, including V1, which was further validated by the absence of CTB-labeled neurons in the dorsal lateral geniculate nucleus (dLGN), where the thalamic neurons that project to V1 are located (Figures 1C-D).

#### Projection specificity across ages

Across all developmental stages examined, the fraction of dual-labeled V1→AL+PM neurons remained consistently low (P3→P6: 1.4 ± 0.3%, n=3; P6→P9: 2.3 ± 0.5%, n=4; P12→P18: 1.3 ± 0.2%, n=3; P60+→P65+: 2.3 ± 0.7%, n=3; mean ± SEM, Figures 1F-G). These data support Model 2, indicating that V1→AL and V1→PM projections are established via early target-specific axon guidance rather than by exuberant, multi-target innervation followed by pruning.^19^

#### Laminar and timing differences between projections

Examining the developmental dynamics of the populations constituting these dedicated V1→HVA channels, we next analyzed the emergence and laminar distribution of V1→AL, V1→PM, and V1→AL+PM CCPNs across postnatal stages. The data revealed stage-specific projection patterns. At P3→P6, V1 layer 5 (L5) neurons predominantly projected to PM, with minimal labeling from other layers or labeling of V1→AL neurons in any layer. By P6→P9, deep-layer V1→PM projections persisted, and a subset of L5 neurons began projecting to AL. At P12→P18, a small number of V1→PM neurons began to emerge in L2/3 upper, while new V1→AL neurons appeared in both upper and lower L2/3 in addition to L5 V1→AL neurons. Finally, the mature adult pattern had emerged by P60+→P65+. The proportion of V1→PM neurons continued to increase selectively in upper L2/3 to become concentrated in L5 and upper L2/3. In contrast, the proportion of V1→AL projections increased selectively in lower L2/3 and decreased in layer 5 to become localized primarily to L2/3 lower (Figures 1H-I).^7,23,24^ These findings support a spatiotemporally regulated developmental program in which V1 CCPNs extend axons in a target-specific and layer-defined manner, rather than through early, widespread multi-target outgrowth.

Lastly, to assess whether the populations constituting these dedicated V1→HVA channels are non-overlapping throughout the course of development, we investigated the possibility that some early PM projecting neurons prune their PM projection and switch to AL. To test this, we performed sequential injections of long-term labeling retrograde tracers: AAVretro-H2B-eGFP into PM at P3 and AAVretro-H2B-mCherry into AL at P35, harvesting at P45 (Figure 1K). This approach was chosen over CTB, which exhibits relatively weak signal intensity after prolonged incubation, particularly when injection volumes are small. We quantified the proportion of double-labeled V1 neurons out of the H2B-eGFP+ (early PM-projecting) population (Figure 1L). If many neurons redirected axons from PM to AL, we would expect higher double-labeling in the sequential group (model 1). However, the proportion of co-labeled neurons in the sequential injection group did not differ significantly from that in animals that received simultaneous injections in both areas at P35 and were harvested at P45 (Figure 1M). This result, together with our CTB tracing data (Figures 1F-I), demonstrates that dedicated V1→AL and V1→PM projections arise from two largely distinct CCPN populations developing in sequence.

### Global axon output from V1 CCPNs follows a spatiotemporally heterogeneous developmental trajectory

While retrograde tracing offers precise information about neurons projecting to specific targets, it is inherently limited by the number of injection sites and distinguishable fluorophores. To overcome these limitations and expand on the target-specific developmental dynamics revealed through our targeted retrograde tracing experiments, we used a glycoprotein-deleted negative-strand RNA Vesicular Stomatitis Virus (VSV) expressing tdTomato (VSVdG-tdTomato) pseudotyped with VSV glycoprotein (VSVG), which efficiently and rapidly labels axons within four days of injection (Figure 2A). Using this rapid anterograde tracing approach, we next mapped the developmental trajectories of V1 CCPN axonal projections to eleven different cortical targets, including HVAs. After harvest, whole brains were coronally sectioned at 50 μm, and were imaged and analyzed using Bell Jar, an image processing software designed for semi-automated axon detection and quantification across brain regions (Figures 2B and S1).^25^ Using Bell Jar, axon projection data were segmented as binary pixel counts and aligned to the Allen Mouse Brain Common Coordinate Framework Reference Atlas (CCFv3) for target areas including RSPv, RSPd, RSPagl, PM (VISpm in CCFv3), AM (VISam), A (VISa), RL (VISrl), AL (VISal), LM (VISl), LI (VISli), and POR (VISpor). We aligned all samples to the adult CCFv3 through an initial step of proportional scaling of the developing brain, followed by manual adjustment of cortical borders based on anatomical landmarks identified from DAPI density and insights derived from marker protein expression patterns (Figure S1).^26–31^ This approach enabled consistent region assignment across developmental stages. To validate Bell Jar’s performance on our dataset, we compared automated annotations to manual annotations by two expert human raters across both densely and sparsely labeled axon images. Bell Jar’s precision and recall scores were nearly indistinguishable from those of human-human comparisons (Figure S2). We also quantified the number of starter neurons (tdTomato+ soma at injection sites) to normalize axon output as axon pixels per starter neuron (Figure 2C).

In the earliest age group (P3→P7), axons preferentially infiltrated medial cortical areas such as RSPv, RSPd, RSPagl, and PM. In contrast, lateral areas including AL, LM, LI, and POR showed relatively sparse axon labeling (Figure 2C), consistent with our CTB data (Figures 1H-I). In the following P12→P16 age group, axonal projections expanded broadly, with increased labeling observed across all target areas (Figure 2F, Tables S2-3). During this phase, medial target areas exhibited relatively reduced proportional axon coverage, while lateral areas such as RL, AL, LM, and LI, displayed dramatic increases (medial area change: 86%→28%, lateral area change: 14%→72%, Figure 2E, top). POR and A showed only modest increases over the same time period, indicating spatially restricted dynamics. By P20→P24, axons were distributed broadly, but specific reductions in axon pixels per starter neuron were observed in lateral areas RL, AL, LM, LI, and POR (Figure 2F, Tables S2-3). These lateral areas exhibited strong increases from P7 to P16, followed by pruning to levels comparable to adult brains. In contrast, medial regions, including A, AM, PM, and all RSP subregions, followed a more gradual pruning trajectory with mature axon projection patterns not fully established until after P24 and reaching adult-like states by P60+→P64+ (Figure 2C-F).

We further analyzed our data to examine the laminar distribution of V1 CCPN axons in the eleven queried cortical areas using depth profiles generated from Bell Jar-processed images (Figures 3 and S4). For each animal, axon signals were collapsed across coronal sections to create 1×100 arrays representing signal intensity from pia mater to white matter. Clustering of these depth profiles (n=139, target areas) using Pearson’s correlation and Ward’s method identified two major projection types: Cluster 1 (deep-layer, DL; n=50) and Cluster 2 (multi-layer, ML; n=89), along with a non-projecting (NP) group where axons were not detected (n=15) (Figure 3B-C). Tracking these clusters across development revealed that medial HVAs were dominated by DL-type projections at P3→P7, while lateral HVAs showed mostly NP profiles, indicating little to no input. By P12→P16, ML projections became prominent in lateral HVAs (Figure 1F). These results suggest that laminar targeting of V1 CCPNs follows distinct, area-specific developmental trajectories.

**Figure 3.**
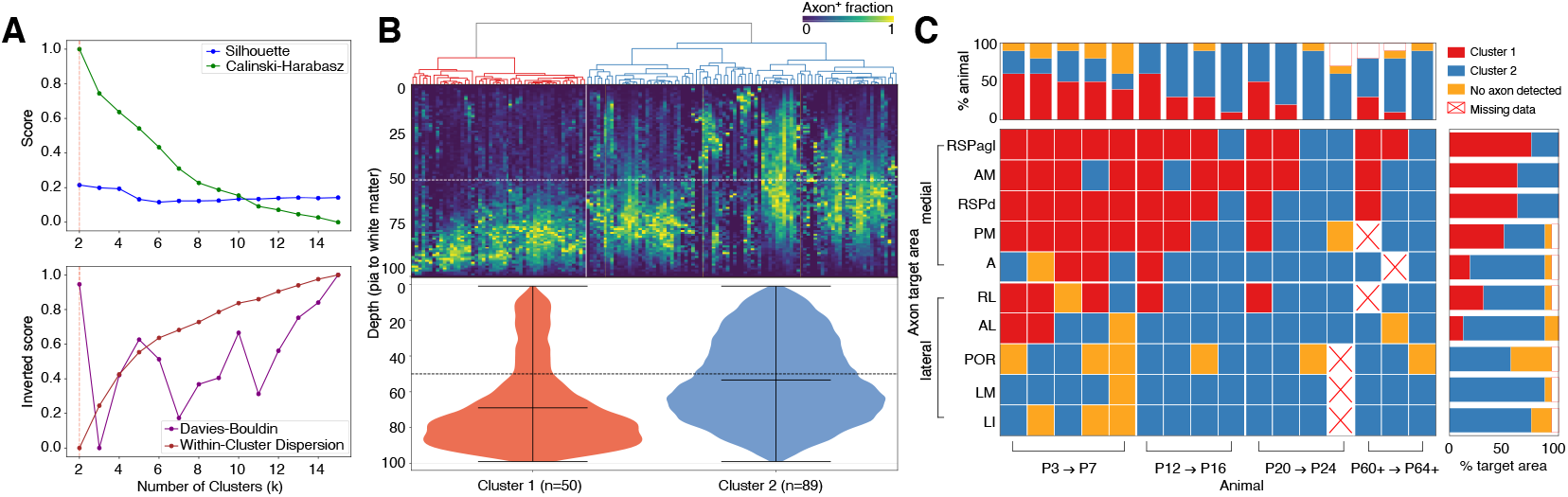
Spatiotemporal dynamics of axon distribution profiles in V1 CCPN target areas across development. **(A)** Evaluation of clustering performance using multiple metrics: Silhouette score, Calinski-Harabasz index, Davies-Bouldin index, and within-cluster dispersion for V1 CCPN target areas. The dotted red lines indicate the selected number of clusters (k = 2). **(B)** Hierarchical clustering of V1 CCPN target areas across developmental and adult stages using the Ward’s method and Pearson’s correlation coefficient as the distance metric, based on axon fraction profiles measured along the pia-to-white matter axis. Violin plots show axon distributions across cortical depth for each cluster. **(C)** Developmental shifts in cluster assignment for 10 cortical areas targeted by V1 CCPNs, revealing spatiotemporal changes in laminar axon profiles.

Our findings show that the V1→PM before V1→AL developmental dynamics, revealed through our targeted retrograde tracing experiments (Figure 1), generalizes to a broader medial-before-lateral scheme for areas beyond these two target areas, PM and AL. Additionally, these findings demonstrate that V1 CCPNs undergo regionally and temporally distinct phases of exuberance and pruning. Rather than following a uniform developmental timeline, axonal refinement proceeds according to a spatiotemporally organized plan: projections to medial HVAs emerge earlier but mature more slowly, while those to lateral HVAs arise later and refine more rapidly. Together, our results reveal a novel, target-dependent developmental program marked by complex and heterogeneous dynamics in the formation of cortico-cortical connectivity.

### Axonal pruning, rather than apoptosis, underlies the reduction of laterally projecting V1 axons during the second postnatal week

Having observed exuberant growth and subsequent reduction of axon coverage in both medial and lateral HVAs in our bulk axon tracing data (Figure 3), we next sought to determine the mechanism accounting for the observed reduction. We hypothesized that this effect could be due to either axonal pruning or apoptosis of V1 cells that extend axons to their target HVAs early on (Figure 4A).^32^ To distinguish between these two possibilities we injected AAVretro-H2B-eGFP into LM at P10 and incubated the virus for either 5 or 20 days (Figure 4B). We chose these time windows as the axon coverage reduction effect is shown to the greatest extent in lateral HVAs between the P12→P16 and P20→P24 age groups. Our injection-incubation strategy was designed to label V1 CCPNs extending axons laterally during the peak of exuberant growth, and then to follow these cells before and after the subsequent reduction in axon coverage. If the observed reduction in axon coverage is caused by apoptosis, fewer retrogradely labeled cells should be present in V1 in long incubation (20 days) animals than in short incubation (5 days) animals. Likewise, if retrogradely labeled V1 cells are not reduced after 20 days of incubation, then we assume it is the axons which are selectively lost rather than the whole cell (Figures 4A-B).

**Figure 4.**
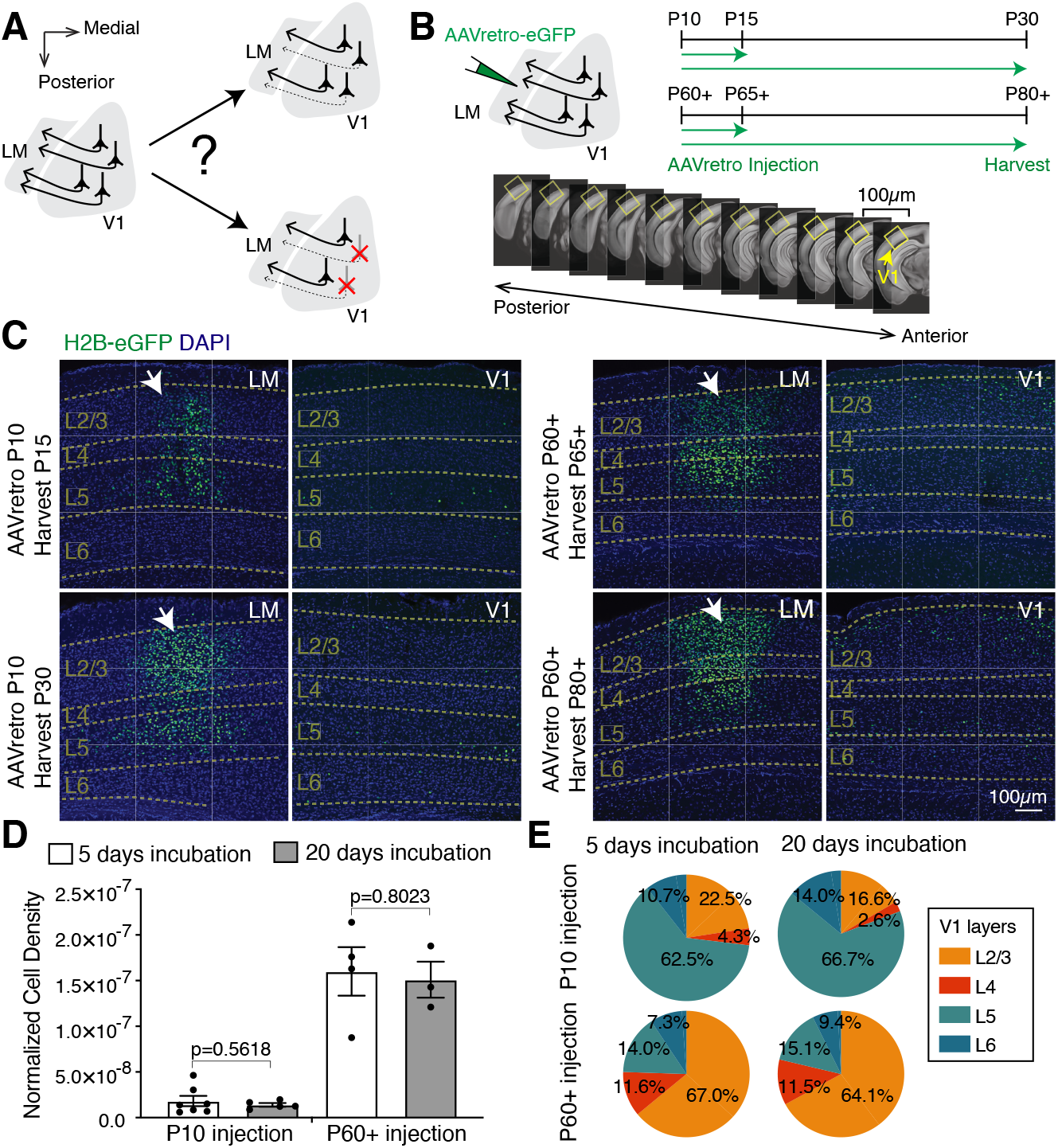
Axonal pruning, rather than apoptosis, underlies the reduction of laterally projecting V1 axons during the second postnatal week. **(A)** Two conceptual models illustrating potential mechanisms of axon reduction in lateral V1 projections: axonal pruning (top) versus neuronal apoptosis (bottom). **(B)** Schematic of the experimental design. Top left: AAVretro-H2B-eGFP was injected into LM to label V1→LM neurons retrogradely. Top right: Timeline of viral injection and brain collection at various developmental and adult stages. Bottom: sequential coronal images across the anterior-posterior axis of the mouse V1 used for whole-region analysis (adapted from the Allen Brain Common Coordinate Framework^27^). **(C)** Representative confocal images showing local H2B-eGFP+ infections (white arrows, left columns) on LM injection sites, along with retrogradely labeled H2B-eGFP+ CCPN nuclei (green) in V1. **(D)** Quantification of V1→LM cell nuclei, normalized to V1 volume, viral labeling efficiency, and brain growth in P30 brains after 20-day viral incubation. **(E)** Pie charts showing the laminar distribution of H2B-eGFP+ CCPN nuclei in V1 for four different experimental groups: P10→P15, P10→P30, P60+→P65+, and P60+→P80+. n = 7 mice for P10→P15, n = 5 for P10→P30, n = 4 for P60+→P65+, and n = 3 for P60+→P80+. Student’s t-test; ns = not significant. Data are presented as mean ± SEM.

We quantified retrogradely labeled V1→LM CCPNs across every other coronal section through the entire V1 (Figure 4B bottom), measured the area of tissue containing labeled cells, and calculated cell density (cells/area) for each section (Figure 4C, Table S4). These values were normalized to the number of labeled cells at the injection site to control for variation in injection efficiency and potential increases in eGFP expression over time, as well as to a growth factor to control for brain size increase between P15 and P30. We found no significant difference in the number of retrogradely labeled V1→LM CCPNs between the 5-day and 20-day incubation groups (p = 0.5618, Student’s t test, Figures 5D and S3D), supporting axon pruning as the mechanism underlying the observed reduction in axon coverage in lateral HVAs between P16 and P24. As a control experiment, a similar analysis in adult P60+ animals also showed no significant difference (p = 0.8023, Student’s t test), suggesting that neither viral toxicity nor expression levels affected our results (Figure 5D). Analysis of the laminar distribution of retrogradely labeled V1→LM CCPNs revealed no differences between short and long incubation groups within either young animals or adults (Figure 5E). However, developmental versus adult comparisons showed a clear shift from predominantly deep layer labeling in pups to increased upper layer representation in adults (Figure 5E), consistent with our CTB tracing results (Figures 1H-I). Taken together, these findings indicate that developmental reduction in lateral axon projections results from selective axon pruning rather than neuron loss (Figure 4A, top right).

**Figure 5.**
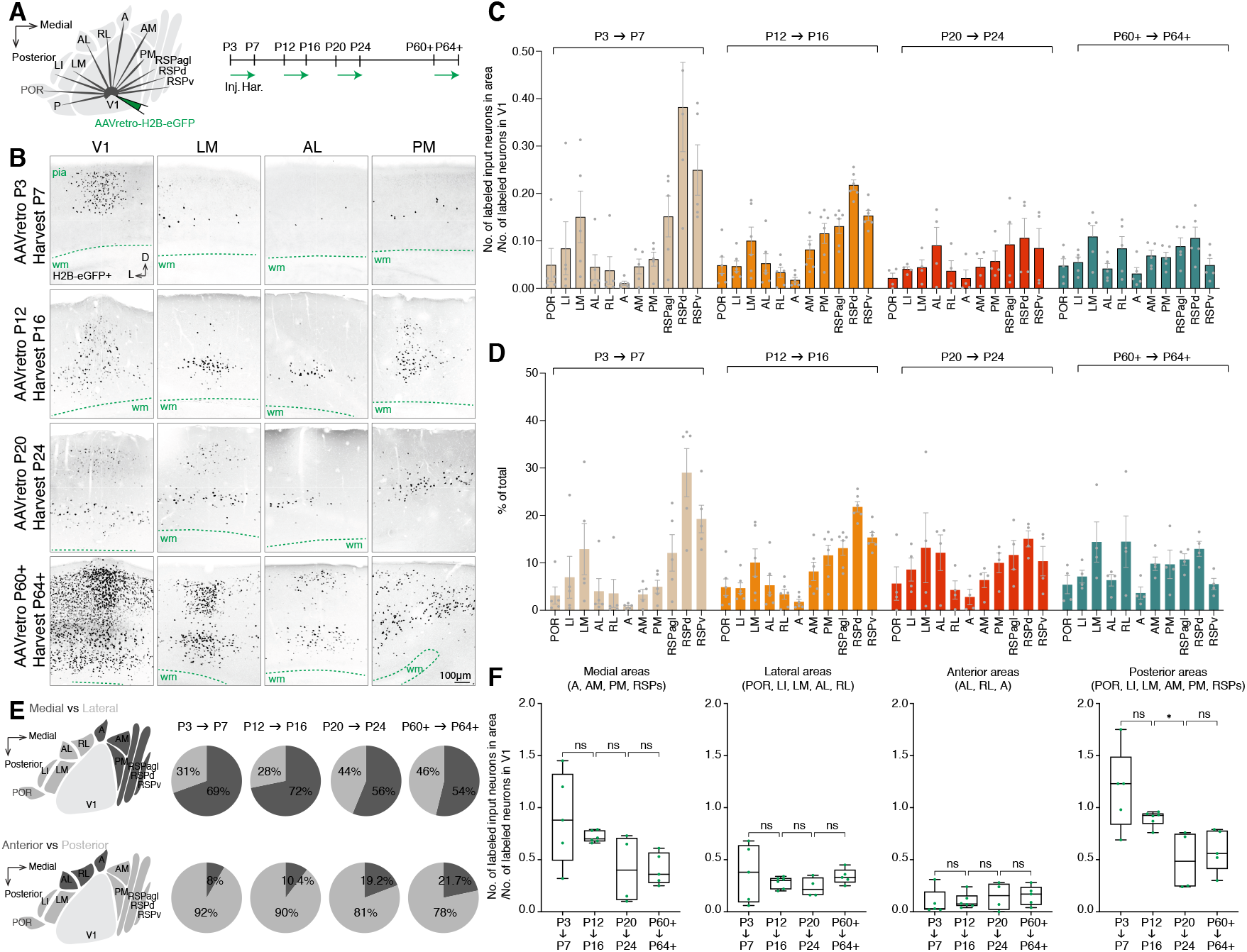
Reconstruction of long-range input neurons from eleven cortical areas to V1 across postnatal stages. **(A)** Experimental design: AAVretro-H2B-eGFP was injected into the center of V1 at various developmental and adult stages. Brains were collected 4 days post-injection for analysis of retrogradely labeled input neurons. **(B)** Representative confocal images of coronal brain sections showing locally infected H2B-eGFP+ neurons on V1 injection sites (leftmost column), along with retrogradely labeled H2B-eGFP+ input neurons in selected HVAs: LM, AL, and PM. **(C-D)** Quantification of input neurons across eleven cortical areas throughout development: **(C)** Number of H2B-eGFP+ neurons in each input area normalized to the number of labeled neurons in V1; **(D)** Percentage of total labeled input neurons attributed to each area. **(E)** Pie charts showing the distribution of H2B-eGFP+ input neurons in medial versus lateral (top) and anterior versus posterior (bottom) input areas at different developmental and adult stages. **(F)** Comparative analyses of normalized input neuron numbers across developmental stages in medial, lateral, anterior, and posterior input area groups. n = 5 mice for P3→P7, n = 6 for P12→P16, n = 4 for P20→P24, and n = 5 for P60+→P64+. one-way ANOVA with uncorrected Fisher’s LSD; ns = not significant, *p < 0.05. Data are presented as mean ± SEM.

### Developmental trajectories of feedback inputs to V1 and emergence of reciprocal cortico-cortical connectivity

To complement our VSVdG-mediated axon output tracing of V1 to eleven cortical areas (Figure 3), we examined the developmental trajectory of feedback input from the same areas to V1 using retrograde labeling with AAVretro-H2B-eGFP (Figure 5A). Injections of AAVretro-H2B-eGFP into V1 were performed across the same postnatal time windows as the bulk anterograde tracing experiments (P3→P7, P12→P16, P20→P24, and P60+→P64+), after which coronal sections (50 μm) were imaged and registered to the Allen CCFv3 atlas using Bell Jar. Retrogradely labeled H2B-eGFP+ feedback input neurons were manually quantified across all eleven higher cortical areas (Figure 5B-C and S3), and counts were normalized to the number of labeled starter neurons at the V1 injection site to account for variability in infection and tracing efficiency.

Based on normalized feedback neuron counts and their proportions across cortical areas, we did not observe statistically significant changes in the overall magnitude or distribution of higher-order cortical inputs to V1 during development, although a mild decreasing trend was evident (Figures 5C, F, Tables S2-3). In contrast to the “medial first” projections from V1, substantial feedback was already present from both medial and lateral areas at P3→P7, indicating that feedback projections from lateral HVAs precede the emergence of feedforward projections to those areas. Furthermore, the medial-lateral and anterior-posterior distribution of feedback input neurons remained largely stable across developmental stages. However, there was a tendency for reduced input from medial areas (RL, AL, LM, LI, POR) relative to lateral areas (A, AM, PM, RSPagl, RSPd, RSPv), as well as from posterior regions (AM, PM, RSPagl, RSPd, RSPv, LM, LI, POR) relative to anterior regions (A, RL, AL) (Figure 5D-E). Notably, we observed an early peak in feedback input from RSPd and RSPv at P7, followed by a marked decline by P16 (Figure 6B). These data suggest that feedback inputs from medial HVAs undergo more dynamic developmental changes compared to those from lateral HVAs.

**Figure 6.**
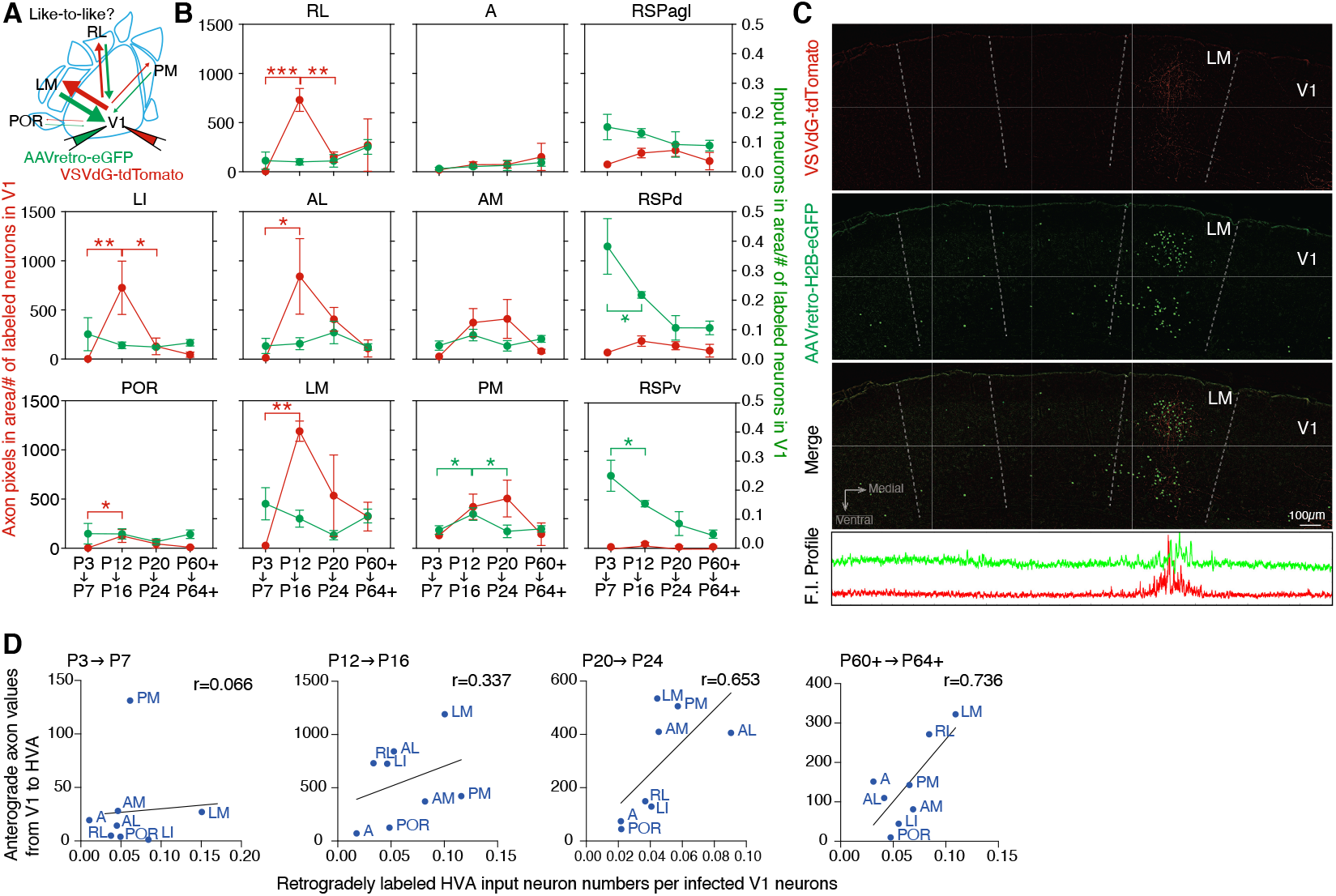
Correlative analysis of connectivity between V1 and HVAs reveals the gradual emergence of like-to-like reciprocal connectivity across development. **(A)** Schematic summarizing the experimental datasets used for correlation analysis between V1→HVAs and HVAs→V1connectivity across development. **(B)** Time course quantification of V1→HVAs axon outputs (red, detected axon px/starter) and bulk HVAs→V1 (green, labeled cells in HVAs/labeled V1 cells at injection site) detected in various target areas at different postnatal stages. One-way ANOVA with uncorrected Fisher’s LSD test. *p < 0.05, **p < 0.01, ***p < 0.001. Data are presented as mean ± SEM. **(C)** Representative confocal images showing colocalization of local V1 axons (tdTomato+) and input neurons (H2B-eGFP+) in LM. Fluorescence intensity profiles of tdTomato and H2B-eGFP signals along the lateral-medial axis of a coronal section (bottom). **(D)** Pearson’s correlation coefficients (r) between V1→HVAs and HVAs→V1 across development. Pearson’s correlation of averages by area, P3→P7: r=0.0655 r^2^=0.0043, P12→P16: r=0.3366 r^2^=0.1133, P20→P24: r=0.6533 r^2^=0.4269, P60+→P64+: r=0.7363 r^2^=0.5421). Sample sizes for V1→HVA axons: n = 5 mice for P3→P7, n = 4 for P12→P16, n = 4 for P20→P24, and n = 3 for P60+→P64+. Sample sizes for HVA→V1 input neurons: n = 5 mice for P3→P7, n = 6 for P12→P16, n = 4 for P20→P24, and n = 5 for P60+→P64+.

To understand how reciprocal cortico-cortical connectivity develops, we compared the trajectories of feedforward projections from V1 to higher visual areas (HVAs) with feedback inputs from HVAs to V1 (Figure 6B). Rather than showing consistent correlation or anti-correlation, feedforward and feedback projections for each V1–HVA pair followed distinct developmental courses, suggesting that these pathways mature independently. Mature interareal connectivity in the mammalian cortex is often reciprocal, with the strength of projections between two areas approximately balanced in both directions.^33,34^ While this organizational feature has been documented at both areal and cellular levels across species,^7,34,35^ its developmental origins remain unclear. In our datasets, regions that received strong V1→HVA projections tended to overlap spatially with those sending feedback to V1, suggesting an early emergence of reciprocity (Figure 6C).

To test whether this reciprocity strengthens over time, we compared the spatial profiles of V1 outputs and HVA inputs across ages using Pearson correlation analysis (Figures 3 and 5). This analysis revealed a progressive increase in correlation, from near zero in early development (P3→P7: r = 0.066, p = 0.8775) to a significant positive correlation in adulthood (P60+→P64+: r = 0.736, p = 0.0373) (Figure 6D). These findings suggest that reciprocal cortico-cortical connectivity between V1 and HVAs is gradually refined during postnatal development rather than specified simultaneously. This refinement may also partially reflect a temporal mismatch between the P3→P7 and P12→P16 age groups, during which feedback connections from both medial and lateral HVAs emerge concurrently, while feedforward projections initially target only medial HVAs.

### MAPseq-based high-throughput mapping of individual V1 CCPNs shows that projection motifs are largely comparable across development at the single-cell level

To study the development of long-range axonal targeting at single-cell resolution, we used MAPseq^22^ to examine feedforward axonal projections during three postnatal windows: P14→P16, P22→P24, and P60+→P62+. Earlier developmental stages were excluded from analysis due to overall low levels of axonal projections (Figure 2C) and challenges in reliably micro-dissecting target areas. While bulk anterograde and retrograde tracing of feedforward and feedback connectivity between V1 and HVAs reveals overall connectivity patterns, it does not resolve how the projections and collaterals of individual neurons are organized. MAPseq overcomes this limitation, as it is a high-throughput barcoding method in which hundreds to thousands of V1 neurons are uniquely labeled following a single injection of an RNA barcoded virus library. Since barcodes are transported along axons, we dissected five known target areas of V1 CCPNs and sequenced the barcodes in each region to identify projection motifs and quantify projection strengths of individual neurons across development.

To profile MAPseq-labeled axons across three postnatal time points, we quantified total unique molecular identifier (UMI) counts, mean UMI counts per cell, and the number of unique barcodes in the five chosen targets. These metrics in P60+→P62+ brain samples were generally consistent with those from the VSV-labeled population dataset (Jensen–Shannon divergence values, 0.2597 for the comparison between MAPseq and VSV datasets, and 0.1510 for the comparison between MAPseq and the Allen Brain Projection Atlas datasets, Figures 2 and S3). LM/LI consistently showed the highest total UMI counts, highest mean UMI counts per barcode, and greatest barcode diversity, while RSP exhibited the lowest levels across these metrics (Figure 7E). Notably, the mean UMI count in LM/LI decreased from P14→P16 to P22→P24 (one-way ANOVA, P=0.0497, Figure 7E), which resembles the decline observed in VSV-labeled axons in LM and LI from P12→P16 to P20→P24 in the VSV tracing experiments (Figure 2C).

**Figure 7.**
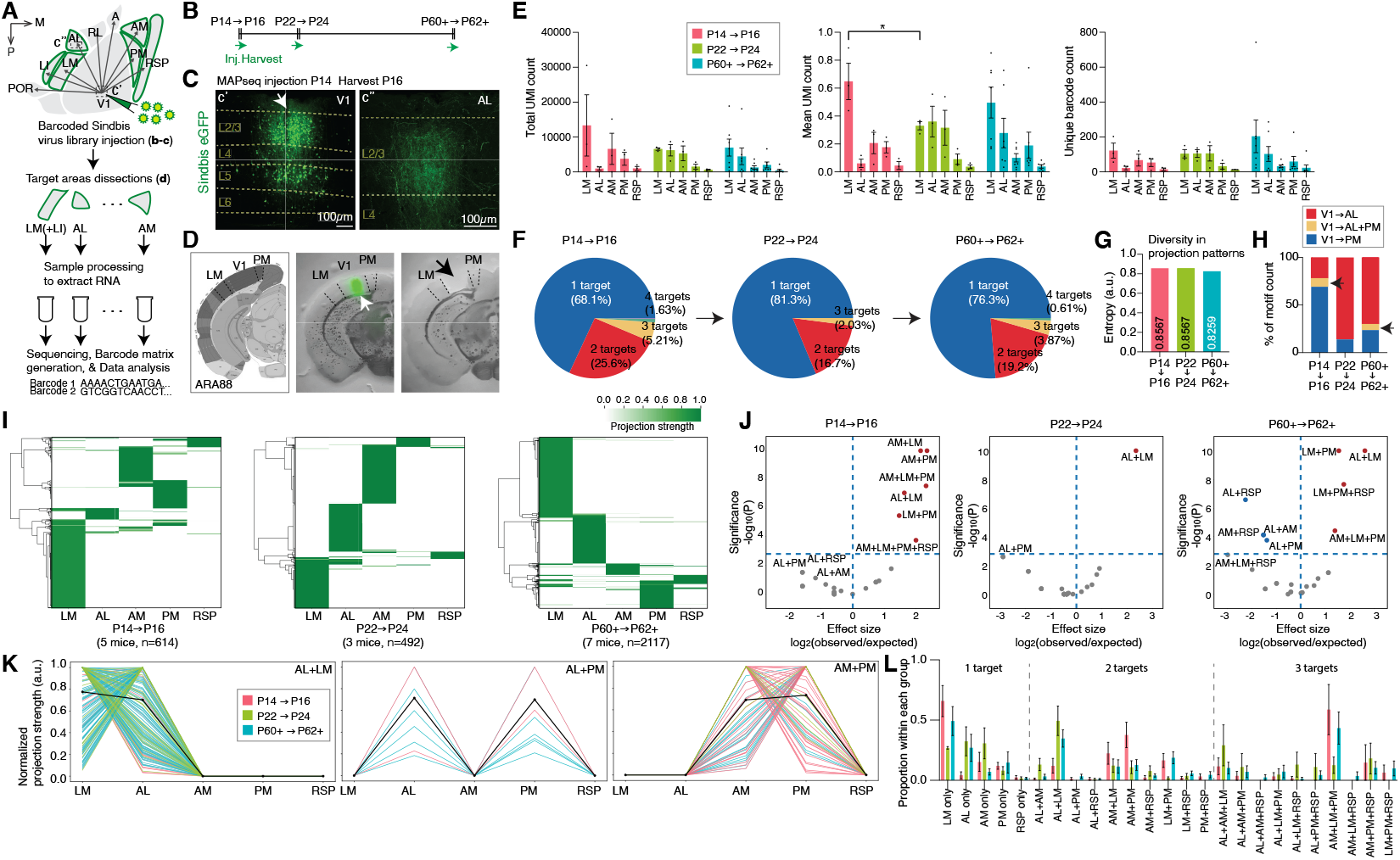
High-throughput, single-neuron resolution analysis of axonal projection pattern development using MAPseq. **(A)** Experimental design for MAPseq. A barcoded Sindbis virus library was injected into the center of V1 at various developmental and adult time points. Brains were collected 48 hours later, and target regions, including LM/LI, AL, AM, PM, RSP, and the cerebellum (as a negative control), were microdissected. Tissue samples were processed for RNA extraction, followed by Illumina sequencing and barcode matrix generation for projection pattern analysis. **(B)** Timeline of viral injection and brain collection at various developmental and adult stages. **(C)** Confocal images showing eGFP+ barcoded Sindbis virus injection into V1 at P14 (arrow, left) and eGFP+ barcoded axons in a representative target region, AL (right). **(D)** Example of a Sindbis eGFP-labeled V1 region (white arrow) from a coronal section before (middle) and after (right) microdissection (black arrow). This section’s cortical areas - LM, V1, and PM - correspond to those of the ARA88 section from the Allen Brain Common Coordinate Framework^27^. **(E)** UMI sum (left), UMI mean (middle), and barcode sum (right) for individual animals across three developmental stages. n = 3 mice for P14→P16, n = 3 for P22→P24, and n = 7 for P60+→P62+. One-way ANOVA with uncorrected Fisher’s LSD. p < 0.05. Data are shown as mean ± SEM. **(F)** Pie charts showing the distribution of neurons projecting to 1–4 targets in each age group (P14→P16, P22→P24, P60+→P62+). **(G)** Diversity of projection patterns measured at different developmental stages. **(H)** Proportion of neurons showing specific projection motifs (V1→AL, V1→PM, V1→AL+PM) across age groups. **(I)** Heatmaps of projection strength from barcoded neurons to five targets, measured by MAPseq at three postnatal stages. **(J)** Volcano plots showing statistical significance and effect sizes for over- and under-represented projection motifs across developmental stages. See Methods for null model generation. n = 614 cells, 5 mice (P14→P16), 492 cells, 3 mice (P22→P24), 2117 cells, 7 mice (P60+→P62+). **(K)** Projection strengths of individual neurons for six representative projection motifs across three developmental time points. See Figure S6 for the complete motifs. **(L)** Proportions of individual motifs within groups of neurons projecting to one, two, or three target groups over development. For both **(K)** and **(L)**, P14→P16 neurons (red), P22→P24 neurons (green), and P60+→P62+ neurons (blue).

In addition, analysis of specific projection motifs (V1→AL, V1→PM, V1→AL+PM) over time revealed that the proportion of V1→AL+PM axons labeled by MAPseq remained consistently low across development (greatest at 8.8% at P14→P16, compared to 6.1% at P60+→P62+), consistent with the low frequency of this motif observed in CTB tracing across developmental and adult stages (Figure 1H). Over time, the proportion of V1→AL projections increased, while V1→PM projections decreased, a trend also seen in the CTB-labeled dataset. In addition, comparison of our P60+→P62+ MAPseq data with a previously published adult dataset revealed similar projection target motifs and strengths (See Discussion/Supplementary Text).^10^ Together, these findings show that our developmental MAPseq data align well with results from other tracing methods (VSV and CTB), and our adult MAPseq data are comparable with existing adult datasets (Figures 1, 2, 7, and S3b).

To determine whether axonal projection profiles to five target areas change over development, we quantified the number of targets innervated by individual V1 neurons at each age. At P14→P16, approximately 68.1% of MAPseq-labeled neurons projected to one target, 25.6% to two, and 5.6% to three. We observed similar distributions at P22→P24 (81.3% for one, 16.6% for two, 2.0% for three) and P60+→P62+ (76.3% for one, 19.2% for two, 3.9% for three), with no significant differences across age groups (one target: p = 0.5535; two targets: p = 0.5535; three targets: p = 0.8189; Kruskal-Wallis test with Dunn’s post hoc test, Figure 7F). These results suggest that the number of projection targets per neuron remains largely consistent over time. Likewise, entropy values, measures of projection complexity, were similar across stages (P14→P16 = 0.8567, P22→P24 = 0.8567, P60+→P62+ = 0.8259) (Figure 7G), further indicating that overall projection patterns are not substantially altered during this developmental window.

Next, we examined the diversity and abundance of individual neurons’ axonal projection motifs and how they change across development. We analyzed 614 MAPseq-labeled V1 CCPNs from 3 mice at P14→P16, 492 neurons from 3 mice at P22→P24, and 2117 neurons from 7 mice at P60+→P62+. To assess motif specificity, we compared the observed data to a null model in which each neuron projects independently to each target with equal probability and no preference for specific motifs. Our analysis revealed a wide range of projection patterns, with certain motifs occurring in a non-random manner in the adult brain, consistent with previous findings (Figures 7J-K and Table S5).^10^ We then asked whether the prevalence of specific motifs changed over time. Similar patterns were observed at P14→P16 and P22→P24, although the statistical significance of individual motifs was generally lower at P22→P24. We also examined whether the relative proportions of projection motifs shifted across ages, but no motif showed statistically significant changes over time (Figure 7L; Kruskal-Wallis test with Bonferroni post hoc correction). These results suggest that specific projection motifs are largely maintained throughout development, with no major shifts in their overall representation. This supports the idea that single-neuron axonal output profiles during development resemble those in the adult brain, rather than being reshaped through extensive inter-regional reorganization, consistent with the results from CTB and sequential AAVretro experiments (Figure 1).

## DISCUSSION

Our study reveals that, despite being spatially intermixed, V1 CCPNs establish long-range cortico-cortical connectivity through axonal development programs that are both target-specific and temporally distinct. These differences include variation in both the onset of axon extension and the timing of pruning, yet the overall projection motifs of individual neurons remain largely stable through postnatal development. This indicates that long-range cortical circuits are refined primarily by local remodeling within target areas, rather than large-scale rewiring of projection identity.

By mapping V1 projections across eleven HVAs during the first three postnatal weeks, our study provides a comprehensive view of long-range circuit formation within a single sensory modality. This aligns with prior studies comparing projection patterns across different sensory and motor systems and highlights the fine spatial and temporal specificity of cortical wiring programs.^19,36^ Notably, we identify differential pruning dynamics along the medial-lateral axis relative to V1, pointing to distinct developmental timelines even within a single sensory system.

The finding that spatially intermixed V1 CCPNs follow divergent developmental paths is notable. While their cell bodies are likely exposed to similar global molecular cues such as morphogens and axon guidance signals, their long-range axons located far from the soma may encounter distinct local cues. The comparable distances from V1 to each HVA further suggest that differences in axonal outgrowth timing are not simply explained by projection length. However, whether features of the white or deep gray matter tracts or surrounding environment differentially affect axonal navigation remains unknown.^37,38^ This suggests that intrinsic factors, such as gene expression profiles, soma positions within the cortical layer, or neurogenesis timing, may contribute to target-specific connectivity programs. Previous work has shown that subsets of CCPNs, for example V1→AL and V1→PM neurons occupy different positions within L2/3 and exhibit gene expression differences, and furthermore sub-laminae of L2/3 exhibit developmental gene expression gradients.^7,24^ Considering the inside-out pattern of cortical development, it will be important to determine whether these subpopulations differ in birthdate and how this might influence their axonal growth trajectory.

In addition to intrinsic programs, our findings raise the possibility that target-derived molecular cues or activity-dependent mechanisms contribute to the temporal regulation of V1 CCPN axonal refinement. The observed differences in pruning timing across visual areas may reflect target region-derived trophic factors or local patterns of neuronal activity.^39^ Interestingly, dorsal stream HVAs (AL, PM, RL, AM) have been reported to exhibit more delayed maturation of visual receptive properties compared to ventral stream areas (LM and LI), largely consistent with our findings demonstrating more rapid attainment of adult-like axon coverage in these ventral stream areas.^40,41^ Binocular enucleation at birth has been shown to reduce V1 to HVA connectivity and the number of HVA targets in the mouse visual cortex.^42^ These findings raise the possibility that differences in the anatomical development of V1–HVA connectivity may be linked to differential functional maturation across HVAs, potentially mediated by activity-dependent interactions between V1 CCPNs and their HVA targets. Moreover, recent whole-brain transcriptomic studies in visually deprived or enucleated mice have shown altered expression of cell adhesion molecules, axon guidance cues, and morphogens, suggesting experience and environmental factors may drive gene regulation–mediated changes in connectivity.^24,43^ Investigating how these molecular signals vary across HVAs and interact with activity patterns during connectivity development will be essential for uncovering the mechanisms that establish target specificity.

Our results also prompt broader questions about how axonal selection and synaptic competition shape circuit refinement. For instance, in regions like LM where some projections are maintained and others are pruned (Figure 3), what determines which axons are stabilized? Is synaptic engagement a prerequisite for retention, as seen in neuromuscular and cerebellar systems? Addressing these questions will require synaptic-resolution approaches, including axonal imaging with synaptic markers or trans-synaptic tracing to identify functionally integrated axons during development.

Surprisingly, HVA→V1 feedback connections emerge by P7, earlier than the previously reported P11.^44^ This likely reflects the stronger soma-localized signal of AAVretro-H2B-eGFP used in our study compared to dextran amine conjugated to Alexa fluor. Our results show that feedforward and feedback V1-HVA connections emerge concurrently, rather than feedforward V1→HVA connections preceding feedback HVA→V1 connections. In contrast to the dynamic postnatal remodeling observed in feedforward V1 CCPN axons projecting to HVAs, we did not detect asynchronous changes in the number or spatial distribution of feedback input neurons from different HVAs to V1. The notable exception was a prominent early peak in input from medial areas (PM and RSPs) at P7, followed by a sharp reduction by P16 (Figures 5, 6). Although it is important to recognize that quantifying labeled axon pixels and counting neuronal somas reflect fundamentally different aspects of connectivity, as individual neurons can vary greatly in the extent of their axonal arborization,^45^ direct comparisons should be interpreted with caution. Nevertheless, it is noteworthy that V1 CCPNs exhibit earlier axonal innervation of medial HVAs (PM and RSPs), which coincides with an initially higher number of feedback input neurons from these same areas (Figures 5, 6). In the adult cortex, L5 neurons projecting from V1 to PM are more abundant compared to those in other layers, suggesting a layer-specific enrichment that may contribute to the early development of reciprocal connections between V1 and medial HVAs. This early maturation of V1–medial HVAs connectivity may support behaviorally relevant functions during early postnatal stages. For instance, interactions between visual cortices and RSPs could be important for sensory-cognitive roles related to pup behaviors such as suckling shortly after eye opening.^46^

While our study is limited by the resolution and cell-type specificity of tools such as MAPseq and VSVdG, we addressed these constraints by incorporating retrograde CTB tracing to examine layer- and target-specific development of V1 projections (Figure 1). Future advances in single-cell or cell-type-specific labeling and high-resolution imaging will enable deeper dissection of how molecular identity, activity, and experience converge to shape individual neuron trajectories.

In summary, we have identified a developmental framework in which V1 CCPNs exhibit stable projection motifs but follow diverse, target-specific temporal programs of axonal growth and pruning. These findings argue against a purely uniform model of cortical wiring and instead support a model of coordinated heterogeneity, where structured connectivity may arise from the interplay of intrinsic and extrinsic developmental cues. This framework not only deepens our understanding of how precise long-range circuits form but also provides a basis for investigating how these processes may be disrupted in neurodevelopmental disorders characterized by aberrant cortical connectivity.

### Supplementary Text

Direct comparison between our P60+→P62+ MAPseq dataset and the previously published adult MAPseq dataset^10^ is limited by differences in the selected target areas. Our study analyzed five regions (LM/LI, AL, AM, PM, and RSP), whereas the previous study included six (LM, LI, AL, AM, PM, and RL). We combined LM and LI into a single dissection due to the absence of a clearly defined anatomical boundary. RL was excluded because it is difficult to identify and dissect reliably in developing brains. Instead, we included RSP, a non-visual area that nonetheless receives substantial input from V1 CCPNs.

Despite these methodological differences, we found both similarities and distinctions between the datasets. Of the five projection motifs that could be directly compared, three were consistently identified as significantly over-or under-represented: AL+LM and LM+PM (corresponding to LM+LI+PM in the previous dataset) were over-represented, while AL+PM was under-represented in both studies. Compared to the earlier dataset, our results showed a higher proportion of single-target motifs and a lower proportion of multi-target motifs. This difference may reflect both the number of target areas included and the presence of RSP, a non-visual target, in our study. Nevertheless, both datasets reveal significant proportions of neurons with either dedicated or broadcasting projection motifs.

## METHODS

### Resource and materials availability

All requests for resources, information, and materials should be directed to the lead contact, Euiseok J. Kim, This study did not generate new biological reagents. Any additional data reported in this paper will be shared by the lead contact upon reasonable request.

### Experimental animals and husbandry

C57BL/6J mice or mouse strains maintained on the C57BL/6J background were used as wild-type (see Table S1). Both male and female mice were used. The specific ages of experimental animals used are described in Table S1. All mice were housed with a 12-hour light and 12-hour dark cycle and *ad libitum* access to food and water. All animal procedures were performed in accordance with the University of California, Santa Cruz animal care and use committee (IACUC)’s regulations.

### Virus preparation

All adeno-associated viruses (AAVs) and VSVdG were produced by the Salk Institute GT3 Viral Core: scAAVretro-hSyn-H2B-eGFP (referred as AAVretro-H2B-eGFP, 1.16X10^13^ GC/ml), scAAVretro-hSyn-H2B-mCherry (referred as AAVretro-H2B-mCherry, 1.21X10^13^ GC/ml), and VSVG+VSVdG-tdTomato (5.34X10^10^ infectious unit/ml). Sindbis viruses were produced by the Cold Spring Harbor Laboratory MAPseq core: Sindbis Virus barcode library expressing eGFP: L2.2 mappnl or vamp2 (∼2.0X10^9^ infectious particles/ml).

### Animal surgery and injections

For all adult surgeries, anesthesia induction was with either 100 mg/kg of ketamine and 10 mg/kg of xylazine cocktail via intraperitoneal injections, or 3% isoflurane and subsequent maintenance with ∼1% isoflurane. Adult mice were mounted in a stereotaxic frame (RWD instruments) for surgery and stereotaxic injections. Stereotaxic injections in adult mice were targeted using the following coordinates: 1.6-1.9 mm rostral, 1.5 mm lateral relative to lambda and 0.4-0.6 mm ventral from the pia for PM, 1.1 mm rostral, 2.6 mm lateral to lambda and 0.4-0.6 mm ventral from the pia for V1, 1.5-2.0 mm rostral, 3.6 mm lateral to lambda and 0.4-0.6 mm ventral from the pia for AL, 0.7-1.0 mm rostral, 3.65 mm lateral relative to lambda and 0.4-0.6 mm ventral from the pia for LM. Injections of all virus types were done using air pressure manually controlled by a 3 ml syringe with 18G tubing adapter and tubing connected directly to the glass injection pipette. To prevent injection backflow, the pipette was left in the brain for 3 minutes after the completion of the injection. Following recovery, all mice were injected with subcutaneous carprofen (5 mg/kg) and housed with a water bottle containing ibuprofen (30 mg/kg).

For all pup surgeries, anesthesia induction was achieved with 3% isoflurane and subsequent maintenance with 0-3% isoflurane. Animals younger than P12 were mounted in a custom restraint frame designed in-house for typical stereotaxic surgeries in pups. Stereotaxic injections in P3 mice were targeted using major blood vessel landmarks and the following coordinates: 1.2 mm rostral from the transverse sinus, 1.0 mm lateral relative to lambda and 0.1-0.25 mm ventral from the pia for PM, 0.7 mm rostral from the transverse sinus, 1.7 mm lateral to lambda and 0.1-0.25 mm ventral from the pia for V1, 1.5 mm rostral from the transverse sinus, 2.45 mm lateral to lambda and 0.1-0.25 mm ventral from the pia for AL, 0.7 mm rostral from the transverse sinus, 2.45 mm lateral relative to lambda and 0.1-0.25 mm ventral from the pia for LM. For ∼P12 and ∼P20 surgeries, injections were guided by major blood vessel landmarks with post-surgery verifications. Accurate targeting of designated areas was screened prior to data analysis by assessing fluorescence at the injection sites and identifying injection locations using brain atlases and the Bell Jar as reference tools.

For cholera toxin subunit B (CTB) tracing experiments, 5–10 nl of CTB conjugated to Alexa Fluor dyes, CTB-647 (1 mg/ml, Invitrogen C34778), CTB-555 (1 mg/ml, Invitrogen C22843), or CTB-488 (1 mg/ml, Invitrogen C34775), was injected into PM and AL, with fluorophore assignment rotated evenly across replicates. Mice were incubated for 3–6 days prior to tissue collection for histology. For VSVG+VSVdG-tdTomato and AAVretro-H2B-eGFP/mCherry experiments, viruses were injected at the previously described coordinates, followed by a defined incubation period prior to tissue collection. Injection details for individual animals are provided in Table S1.

### Tissue collection and histological processing

Tissue was harvested following a defined incubation period (see Table S1) using transcardial perfusion with 1X phosphate-buffered saline (PBS) containing heparin (10 U/ml, Sigma-Aldrich H3393), followed by fixation in 4% paraformaldehyde (PFA). Adult (from P60 to P90) or P20+ brains were immediately dissected from the skull and post-fixed in 2% PFA and 15% sucrose at 4°C overnight. Pup (from P3 to P20) heads were post-fixed in 2% PFA and 15% sucrose at 4°C overnight, after which brains were dissected from the skull and post-fixed again in 2% PFA and 15% sucrose at 4°C overnight.

Following post-fixation, both adult and pup brains were cryoprotected by immersion in 30% sucrose in PBS at 4°C until the tissue sank. Using a freezing microtome (Espredia HM430), brains were embedded in 30% sucrose and coronally sectioned at 50 μm. Sections were collected in cryogen for cryoprotection, with half stored at −20°C and the other half washed in 1X PBS with 0.01% sodium azide for further staining and quantification.

Brains injected with CTB or VSVG+VSVdG-tdTomato were sectioned coronally, counterstained with DAPI (4′,6-diamidino-2-phenylindole; ThermoFisher Scientific Invitrogen D1306), and serially mounted onto glass slides using polyvinyl alcohol mounting medium containing DABCO (PVA-DABCO). Slides were allowed to air-dry, then cleaned and stored at 4°C. For a subset of AAVretro-H2B-eGFP/mCherry-injected brains (see Table S1 for animal details), free-floating sections underwent immunohistochemistry to amplify endogenous fluorescence. Sections were incubated overnight at 4°C with goat anti-GFP (1:1000; Rockland 600-101-215, RRID: AB218182) and rabbit anti-dsRed (1:500; Clontech 632496, RRID: AB10013483) primary antibodies in PBS containing 0.5% normal donkey serum and 0.1% Triton X-100. After three 10-minute PBS washes, sections were incubated with Alexa Fluor-conjugated secondary antibodies: Alexa 488 (1:500; Invitrogen A-11055, RRID: AB2534102) or Alexa 568 (1:500; Invitrogen A-10042, RRID: AB2534017). Stained sections were mounted onto gelatin subbed slides, air-dried overnight in the dark, cleared in xylene, and cover slipped using Krystalon mounting medium (Sigma-Aldrich 64969). Prepared slides were cleaned and stored at 4°C.

### Image acquisition

Tissue sections were imaged using a Zeiss AxioImager Z2 widefield microscope equipped with a 10×/0.45 NA objective (working distance (wd): 2.0 mm). High-resolution tiled images were acquired for each section across relevant fluorophore channels and z-depths, typically using 9 focal planes at 5.5 μm intervals. Scan parameters were optimized per experimental group to maximize signal detection, with consistent settings maintained within each group. For datasets requiring cell counting, imaging parameters were adjusted per animal to facilitate accurate identification. For comparative feature analyses, fixed imaging settings were applied across all samples. Selected representative images were acquired using a Zeiss LSM 880 confocal microscope with either a 10×/0.45 NA (wd: 2.0 mm) or a 40×/0.95 NA corr (wd: 0.25 mm) objective. Whole-brain overviews were imaged using an Olympus SZX16 stereomicroscope.

### Image feature analysis for CTB signal quantification

High-resolution images of DAPI-stained coronal sections were used to manually delineate brain regions. An expert anatomist evaluated each image for distinct structural and cytoarchitectural landmarks, and cortical areas (V1, AL, PM, LM, and RSP) along with their laminar boundaries were outlined individually. Cortical laminae were divided into L1, L2/3 upper, L2/3 lower, L4, L5, L6a, and L6b; some layers (e.g., L2/3 subdivisions or L6 and L6a) were merged if necessary. Injection accuracy was assessed after anatomical annotation, and any samples not meeting criteria were excluded. CTB-labeled neurons were manually counted across all layers and channels, including co-labeled cells, by a single rater to maintain consistency.

### Image processing with Bell Jar

Following acquisition, raw fluorescence and DAPI counterstain images were exported as .TIFF files using ZEN 3.1 (blue edition), then maximum projected, reoriented as necessary (rotated/mirrored), and converted to 8-bit grayscale. Each dataset was aligned to the Allen CCFv3 reference atlas using the Bell Jar pipeline, with manual verification and correction of alignment for each section.^25^ Axon segmentation and signal quantification were performed using the integrated Bell Jar–RSAT pipeline. AAVretro-H2B-eGFP+ nuclei were detected using Bell Jar’s default model (tile size = 320, confidence threshold = 0.1, area cutoff = 100).

### Clustering of axon detection mask data

Clustering was performed on the global depth profile dataset, replicating the methods from a previous study.^34^ Pairwise distances (*d*_*x,y*_)were computed for the profiles such that:

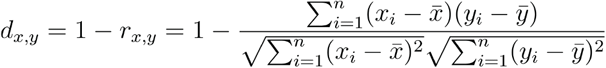

where *r*_*x,y*_ is the Pearson correlation coefficient, *x*_*i*_ and *y*_*i*_ is the *i*^th^ element of the *x* and *y* depth profiles respectively, and 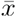 and 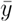 are the sample means of *x* and *y*.

Agglomerative hierarchical clustering was applied via SciPy’s linkage function using Ward’s method. The objective function minimized herein is given by:

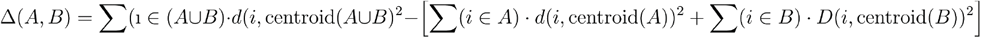

where *A* and *B* are the clusters to be considered for merging and *d*(*i*,centroid(C)) is the correlation-based distance between item i and the centroid of a generic cluster *C* which represents the increase in the sum of squared distances when clusters *A* and *B* are merged. This is consistent with Ward’s assertion in his original article that the generalized agglomerative hierarchical clustering can use any objective function to minimize, wherein here, the above Δ (*A,B*) is used where *d*(*i*,centroid(C)) is correlation-based distance between item i and the centroid of cluster *C*.^47^ The resulting clusters were then sorted by mean depth for purposes of visualization.

Thorough analysis of optimal number of clusters was applied for number of clusters 2 through 15 using Silhouette Score^48^ (k=2), Calinski-Harabasz Score^49^ (k=2), Davies-Bouldin Score^50^ (k=2), Within-Cluster Dispersion score^51^, where k is the optimum number of clusters for each metric. Min-max normalization was applied to the depth profiles just prior to visualization with respect to each profile individually to better represent the relative distributions. Though normalization was not applied prior to clustering, it should be noted that Pearson’s correlation coefficient (the metric used for quantifying distance) is invariant to such transformations since, for random variables *X, Y*,*X* ′ = *a* + *bX*, and *Y* = *c* + *dY* ′:

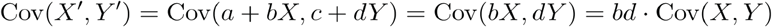

And

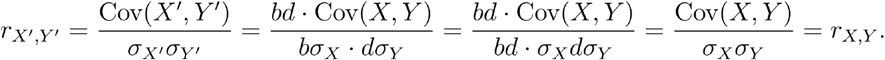

Thus, such normalization offers a representation which is both more intuitive to interpret and consistent with previous literature as well as is mathematically valid (see Figure 3A).

### MAPseq preparation and analysis

#### Tissue preparation and microdissection strategy

A total of 15–100 nl of barcoded Sindbis virus (Cold Spring Harbor Laboratory MAPseq core), depending on the animal’s age (see Table S1), was injected into V1 at postnatal days P14, P22, or P60+ following the procedures described in the ‘Animal surgery and injections’ section. According to MAPseq protocols, brains were harvested 48 hours post-injection, rapidly fresh frozen, and stored at –80 °C until microdissection.

Microdissections were performed under an Olympus SZX16 fluorescent dissecting microscope at a stable temperature of −20°C using an aluminum dissection block semi-submerged in a CaCl_2_ solution containing dry ice. Sections were compared to the Allen CCFv3 by an expert rater and dissected by hand for each region with a scalpel. Collected tissue portions were combined into sample tubes per cortical area from the serial slice dissections.

A secondary microdissection method was developed and used to collect tissue samples from some P60+ brains. Following the fresh harvest of experimental brain tissue samples, specimens were flash frozen to an OCT block and stored at −80°C A specialized reticle was made for an Olympus SZX16 fluorescent dissecting microscope configured for fluorescent dissection. This reticle was a printout of the CCFv3 area borders for all visual cortical areas and retrosplenial cortex as visualized tangential to the posterior aspect of V1 where the viral injections were targeted. This reticle was scaled appropriately using the Olympus SZX16’s magnification adjustment and centered on V1 using common blood vessel landmarks as well as the fluorescent signal at the injection site. The reticle was further aligned to the midline and the transverse sinus. Tissue punches (Ted Pella, 15112-25/50/10) of diameter 0.25 mm, 0.5 mm, or 1.0 mm were then depth-marked for average cortex thickness and used to collect samples from each cortical area. A clean punch was used for every collection site. Following punched tissue collection, the remaining brain tissue was submerged in 8% PFA and allowed to fix overnight. Cryosectioning was then performed on a freezing microtome at 250 μm per section. The resultant tissue was then DAPI-counterstained and mounted to slides for low magnification imaging and post-hoc analysis of the tissue collection.

Microdissected tissues were preserved in TRIzol reagent (ThermoFisher Scientific, 10296-010) and submitted to the Cold Spring Harbor Laboratory MAPseq core facility for downstream processing. RNA extraction, barcoded cDNA library construction, and sequencing were performed using NextSeq High Output PE36 or NovaSeq platforms, following previously established protocols.^22^

#### Barcode processing and projection mapping

Neuronal projection patterns were analyzed at single-cell resolution using a custom Python-based pipeline (Kim laboratory github, https://github.com/Kim-Neuroscience-Lab/mapseq_processing_kimlab). Barcode sequencing data were processed into a matrix where rows represented individual neurons and columns corresponded to anatomical regions (Zador laboratory github, https://github.com/ZadorLaboratory/mapseq-processing).

#### Preprocessing individual barcode matrices *[preprocess_and_aggregate*.*py]*

Values from the negative control, on a per replicate basis, were averaged and the standard deviation was calculated. A minimum threshold filter was applied to each replicate spike in the normalized barcode matrix in a global fashion using this calculated value (AVGneg+STDdev). Subsequently, all cells containing a non-zero value in their negative control sample were removed. As in previous studies, all age-group specific replicate data was then concatenated into a single barcode matrix for further cohort-level analysis.

#### Analysis of cohort level MAPseq datasets *[process-nbcm-tsv*.*py]*

Stringent filtering was applied to the aggregate cohort level data to maximize analysis of “high confidence” cells. This included a requirement that all target counts be less than their V1 count and that the minimum barcode number at the injection site (*UMIinj* ≥ 300 *UMI*), a minimum target barcode count in at least one projection area (UMItarget ≥ 10 *UMI*), and at least a ten-fold difference between the UMIinj and the largest UMItarget on a per cell basis ((UMIinj) ≥ (10(MAX(UMItarget))). Barcodes failing to meet these specifications were eliminated from further analysis.

#### Projection motif identification

Filtered data were used to construct projection profiles by normalizing barcode counts across target regions. Projection motifs were defined as neurons targeting multiple brain regions, with the total number of projections per neuron computed as:

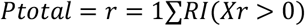

Where *Xr* represents barcode counts in region r_i_, and *I*(*Xr* > 0) is an indicator function.

#### Statistical inference of projection distributions

To estimate the total number of labeled neurons (N_0_), let:

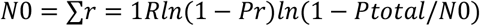

where *Pr* is the probability of a neuron projecting to region r. The probability of observing each projection motif was modeled as:

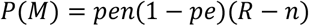

where n is the number of target regions, and (Pe→pe) represents the probability of a neuron forming a projection to e and pe.

#### Cluster and motif analysis

K-means clustering was applied to identify distinct projection patterns, with the optimal cluster number determined via the elbow method. Motif occurrence was compared against an expected distribution, with statistical significance assessed using binomial tests:

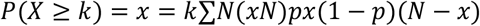

where k is the observed count of a given motif, and p is the expected probability of occurrence. Correction for multiple comparisons was performed using Bonferroni-adjusted thresholds.

Co-targeting probabilities between regions were computed as:

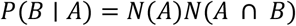

where *N*(*A* ∩ *B*) is the number of neurons projecting to both regions, and *N*(*A*) is the total number projecting to region A.

#### Comparison of VSV and MAPseq data with Allen Brain Institute connectivity data

Adult mouse brain connectivity data were obtained from the Allen Brain Institute (ABI) connectivity atlas (IDs: 100141219, 100147853, 277616630, 277712166, 277713580, 277714322, 304564721, 304585910, 304586645, 304762965, 307137980, 307320960, 307557934, 307593747, 309003780, and 309113907). In these datasets, anterograde axon tracing was conducted by stereotaxic injection of AAV1 expressing eGFP into the right hemisphere’s V1 region, using iontophoresis.^52^ The injection hemisphere for each of the mice used for this comparison was verified visually via the Allen Brain Atlas data portal and the data for each mouse was downloaded as an XML file directly from the portal. The ABI data was filtered to only the relevant areas and its format standardized for analytic ease.

The VSV quantification and MAPseq data for our adult mice were similarly format-standardized to match with the ABI data. For VSV data specifically, the quotient of raw pixels over raw starters was used for the value for each area. For each animal, the areas were assigned the order POR, LI, LM, AL, RL, A, AM, PM, RSPagl, RSPd, RSPv if comparing ABI data to VSV data, and LM+LI, AL, AM, PM, RSP (RSPagl+RSPd) if comparing VSV or ABI data with MAPseq data. When performing the latter, LM and LI as well as RSPagl and RSPd, respectively, we summed together. Standardizing the order (lateral to medial areas) allowed for the data to be compared across datasets. For each dataset, the mean was taken from the subset of values for each area such that each dataset was represented as a single array where each element represents the mean value for that area across all animals.

To normalize the data for comparison across projection patterns, we applied the log-Softmax transformation to each array.^53^ This approach first applies the Softmax function, which converts raw values into a probability distribution using the formula:

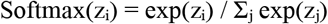

This formulation is mathematically equivalent to the Boltzmann distribution when z_i_ = −ε_i_ / kT, where ε_i_ represents energy states, k is the Boltzmann constant, and T is temperature:

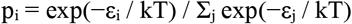

To counteract the tendency of Softmax to flatten small values during exponentiation, resulting in overly uniform distributions, we applied the logarithm of the Softmax output, known as the log-Softmax transformation:

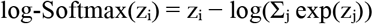

This transformation preserves the relative shape of the original distribution and mitigates artificial inflation of small values.

Following normalization, we calculated pairwise Jensen–Shannon divergence (JSD) between datasets. JSD is a symmetric and bounded measure of similarity between two probability distributions, ranging from 0 (identical distributions) to 1 (maximal divergence). It is defined as:

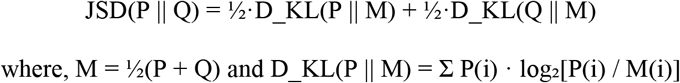

Because interpretation of JSD values depends on the specific context, and no universal thresholds exist, we generated a reference distribution for each dataset by computing all pairwise JSD values among individual samples (resulting in n^2^ values per dataset).^54^ This analysis was performed on projection arrays corresponding to LM+LI, AL, AM, PM, and RSP. For the ABI: n = 16, mean = 0.2749, median = 0.2601, SD = 0.1150, min = 0.0584, max = 0.5492, for VSV: n = 3, mean = 0.4697, median = 0.5218, SD = 0.1122, min = 0.3139, max = 0.5736, and for MAPseq: n = 3, mean = 0.4228, median = 0.4580, SD = 0.1323, min = 0.2018, max = 0.6083. The mean JSD for the ABI set was found to be 0.2749 which indicates that this is how different on average any given array for a pair of mice are in the ABI set. When comparing each set, we have that ABI-VSV = 0.2553, ABI-MAPseq: 0.1510, and VSV-MAPseq: 0.2597 which are all below the mean JSD for the within ABI set.

Kolmogorov-Smirnov p-values were also computed for each pair of datasets for each area. The values for each pair and each area are as follows. For the comparisons between ABI and VSV data: LM+LI = 0.04128, AL = 0.3591, AM = 0.002064, PM = 0.8142, RSP = 0.5046. For the comparisons between ABI and MAPseq data: LM+LI = 0.2492, AL = 0.6504, AM = 0.2492, PM = 0.2054, RSP = 0.001085. For the comparisons between VSV and MAPseq data: LM+LI = 0.4000, AL = 0.8500, AM = 0.1667, PM = 0.9333, RSP = 0.4000.

#### Quantification and statistical analysis

The values of n and what n represents are reported in the Results section or the figure legends. Statistical analyses were performed using GraphPad Prism 10.1.1 or Python 3.9.2. Statistical analyses were selected based on experimental design and included Student’s t-test, chi-square test, one-way ANOVA with uncorrected Fisher’s LSD, two-way ANOVA with Tukey’s multiple comparisons test, Pearson’s correlation, Kruskal-Wallis test with Dunn’s post hoc test, among others. The specific tests used are indicated in the Results section or figure legends. Unless otherwise noted, statistical significance is indicated as follows: ns= not significant, *p < 0.05; **p < 0.01; ***p < 0.001; ****p < 0.0001.

## ACKNOWLEDGEMENTS

We thank Edward Callaway, Bin Chen, David Feldheim, and Yi Zuo for reading the manuscript and Efrain Hernandez-Alvarez for mouse husbandry and histology. We thank Huiqing Zhan and John Hover at the Cold Spring Harbor Laboratory MAPseq core for their technical support and guidance. We thank John Naughton and members at the Salk Institute GT3 virus core for AAVs and VSV reagents. We acknowledge technical support from Benjamin Abrams, UCSC Life Sciences Microscopy Center, RRID: SCR_021135. Purchase of the Zeiss 880 confocal microscope used in this research was made possible through the National Institutes of Health s10 Grant 1S10OD23528-01. We acknowledge support from the UCSC start-up fund, the Whitehall Foundation, the Hellman Fellows Program, the E. Matilda Ziegler Foundation for the Blind, BRAIN Initiative at the National Institutes of Health RF1MH132591, the National Institute of Neurological Disorders and Stroke at the National Institutes of Health R01NS128771 (E.J.K.), and the Koret scholar program (A.L.R.S.).

## Code availability

All code developed in the Kim Laboratory and used in this manuscript is available on our Github project pages, which can be found on the lab website (www.ejkimlab.com/code).

## Author contributions

M.W.J. and E.J.K. conceived and designed the study. M.W.J. performed or supervised CTB, VSV, AAVretro-H2B-eGFP, and MAPseq tracing experiments and final analyses for all datasets. J.M.R. performed CTB, MAPseq, AAVretro-H2B-eGFP and sequential AAVretro-H2B-eGFP and AAVretro-H2B-mCherry surgeries, histology, and microscope imaging and selected analyses in collaboration with M.W.J. A.L.R.S. and A.M.M. developed image and data analysis pipelines under direction of M.W.J. J.A.N., H.T.J., and J.A.G.E. performed histology and microscope imaging, and assisted with image analysis under direction of M.W.J. M.W.J., J.M.R., and E.J.K. wrote the manuscript with input from all authors. E.J.K. provided funding and supervised all aspects of the project.

## Declaration of generative AI and AI-assisted technologies in the writing process

During the preparation of this work, the authors used ChatGPT (GPT-4 Turbo, OpenAI) in order to improve readability and language of portions of the manuscript. After using this tool, the authors reviewed and edited the content as needed and take full responsibility for the content of the published article.

## SUPPLEMENTAL INFORMATION TITLES AND LEGENDS SUPPLEMENTAL FIGURES

**Figure S1.**
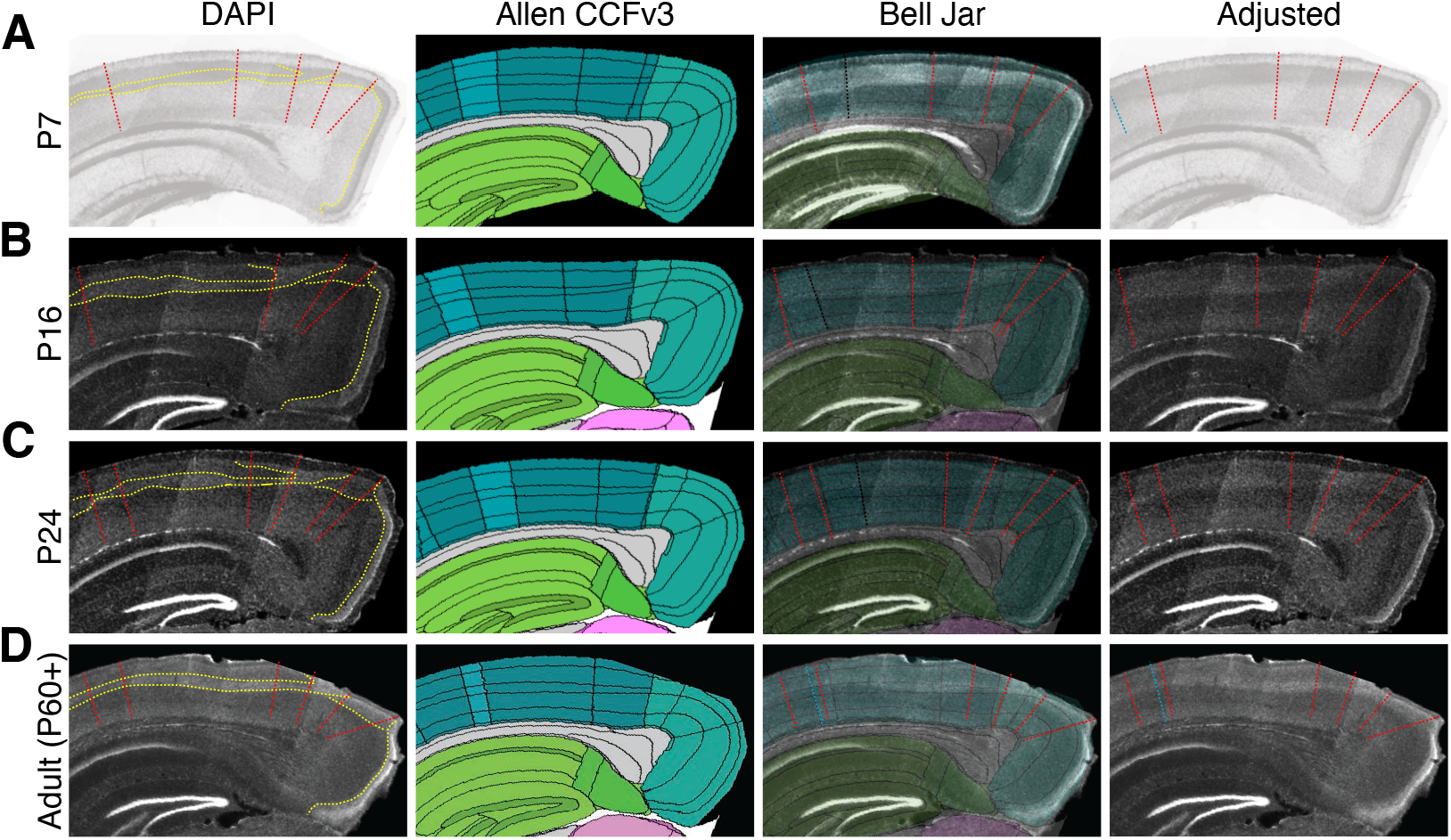
Cortical area alignment and registration across developmental and adult stages using Bell Jar and the Allen Brain CCFv3, related to Figures 1-6. **(A-D)** Representative coronal sections from P7 **(A)**, P16 **(B)**, P24 **(C)**, and adult **(D)** brains showing DAPI-stained visual cortical areas (first column). Yellow dotted lines indicate layer 4 and other anatomical landmarks used to guide area boundary delineation, marked by red dotted lines. Corresponding Allen Brain CCFv3 reference images are shown in the second column. Bell Jar–aligned images are shown in the third column, with newly generated area boundaries (blue dotted lines) and boundaries excluded based on human evaluation (black dotted lines). Final adjusted boundaries (red and blue lines) reflecting manual refinement across developmental stages are shown in the fourth column.

**Figure S2.**
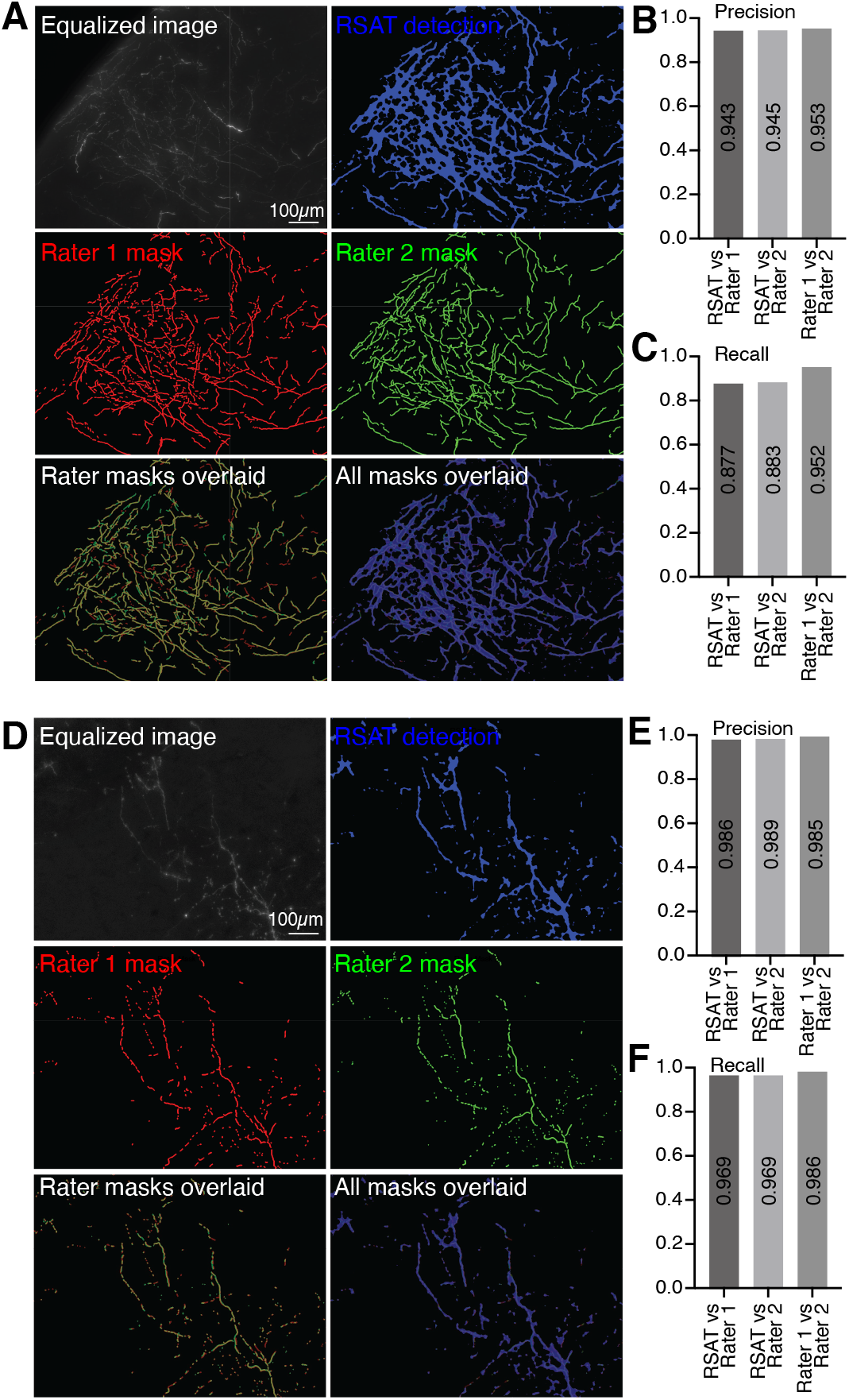
Validation of the Bell Jar-RSAT automated axon quantification pipeline in comparison to expert human raters, related to Figure 2. **(A, D)** Representative fluorescence images showing dense **(A)** and sparse **(D)** tdTomato-labeled axons, displayed as equalized raw images alongside corresponding RSAT automated detection masks and manual annotations by Rater 1 and Rater 2. **(B-C, E-F)** Precision and recall metrics comparing RSAT performance to that of Rater 1 and Rater 2 across both dense and sparse axon labeling conditions.

**Figure S3.**
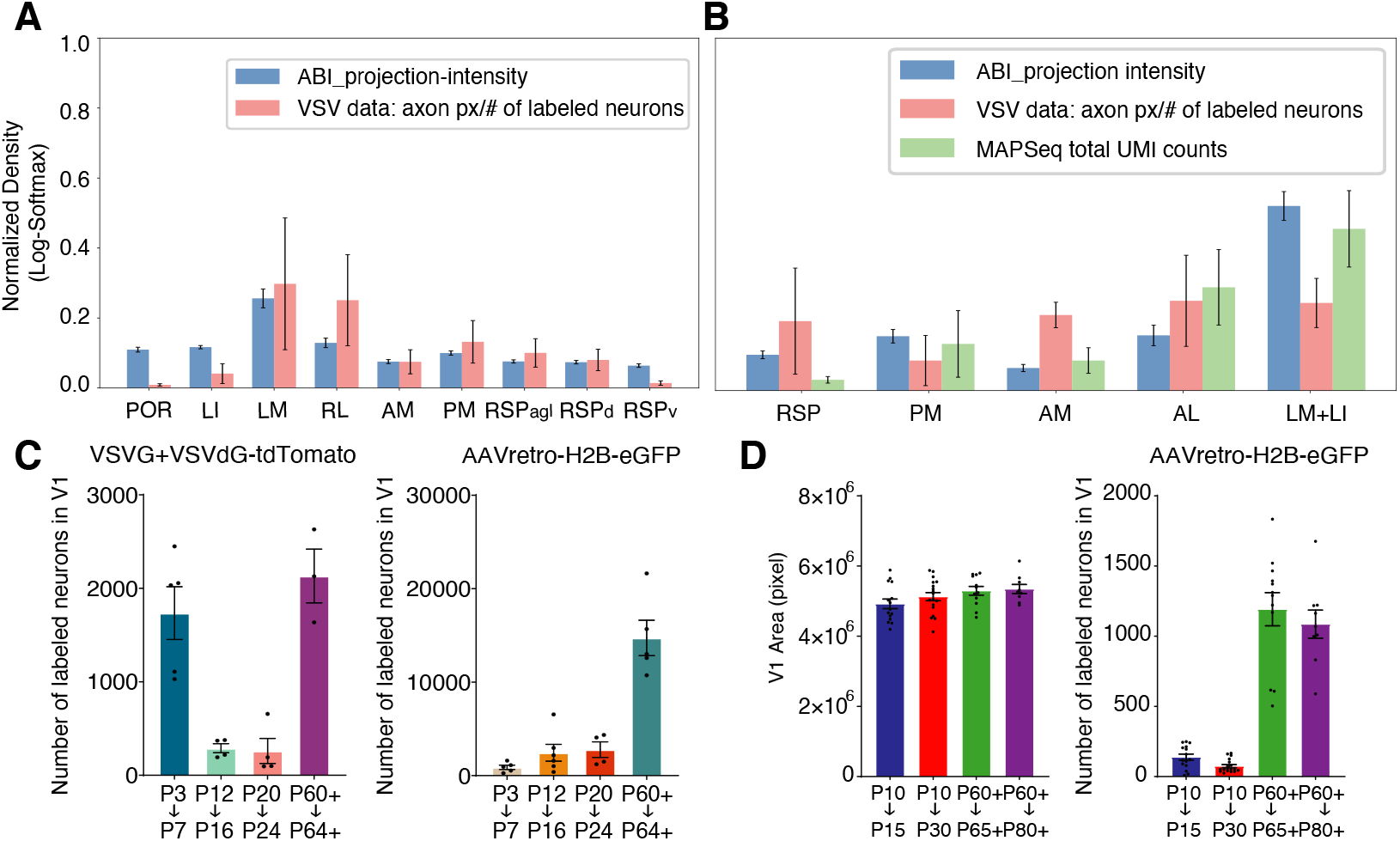
Adult V1 connectivity across tracing methods and developmental quantification of starter neurons, related to Figures 2,4,5, and 7. **(A)** Comparison of adult V1 connectivity data from the Allen Brain Institute (ABI) atlas with VSVG+VSVdG-tdTomato quantification from the P60+→P64 group across multiple HVAs. **(B)** Comparison of three datasets across five micro-dissected areas used for MAPseq analysis: adult V1 connectivity from the ABI atlas, VSVG+VSVdG-tdTomato data from the P60+→P64 group, and total UMI counts from MAPseq in the P60+→P62 group. n=16 mice from ABI connectivity atlas, n= 3 for VSVG+VSVdG-tdTomato, n=7 for MAPseq datasets. Data are presented as mean ± SD. **(C)** Quantification of the number of starter neurons labeled in V1 for both VSVG+VSVdG-tdTomato and AAVretro-H2B-eGFP tracing experiments. For VSVG+VSVdG-tdTomato: n = 5 mice for P3→P7, n = 4 for P12→P16, n = 4 for P20→P24, and n = 3 for P60+→P64+. For AAVretro-H2B-eGFP: n = 5 mice for P3→P7, n = 6 for P12→P16, n = 4 for P20→P24, and n = 5 for P60+→P64+. Data are presented as mean ± SEM. **(D)** Quantification of V1 area size and the number of labeled V1 neurons per section from AAVretro-H2B-eGFP tracing experiments. Data were collected from 15 sections for P10→P15, 18 sections for P10→P30, 12 sections for P60+→P65+, and 9 sections for P60+→P80+, with 3 sections analyzed per animal.

**Figure S4.**
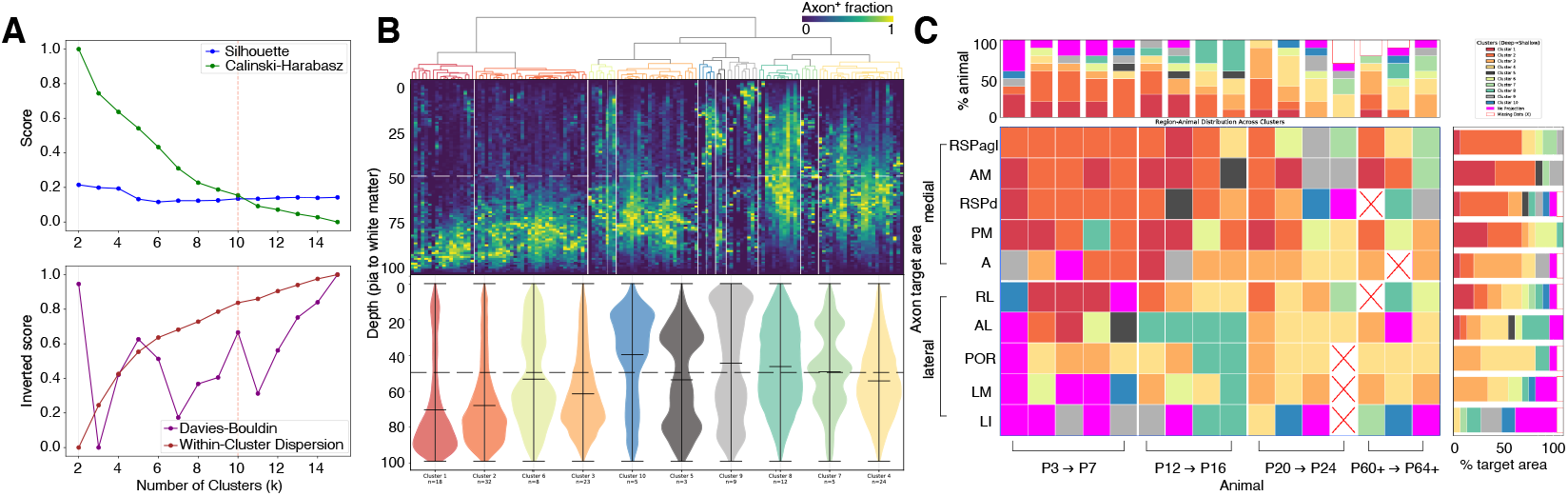
Spatiotemporal dynamics of laminar axon distribution profiles in V1 CCPN target areas across development (k = 10 clustering), related to Figure 3. **(A)** Clustering performance was evaluated using multiple metrics, including Silhouette score, Calinski-Harabasz index, Davies-Bouldin index, and within-cluster dispersion. The red dotted line marks the selected number of clusters (*k* = 10), providing a finer-grained classification than the main figure 3. **(B)** Hierarchical clustering of V1 CCPN target areas across developmental and adult stages using Ward’s method and Pearson correlation as the distance metric, based on axon fraction profiles along the pia-to-white matter axis. Violin plots display the distribution of axonal signals across cortical depth for each of the 10 clusters. **(C)** Developmental transitions in cluster assignments for 10 cortical target areas, illustrating dynamic shifts in laminar axon distribution profiles over time.

**Figure S5.**
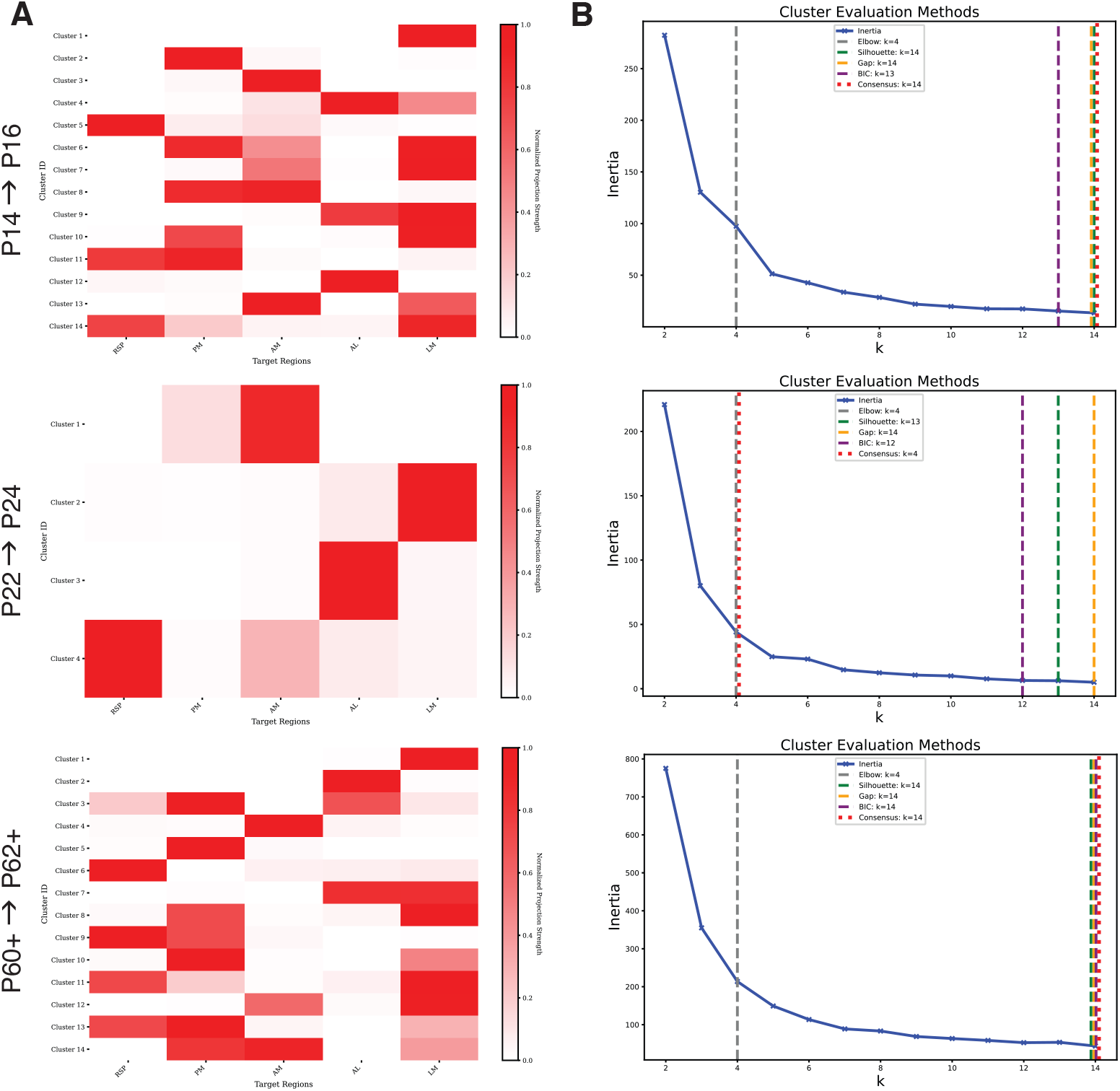
MAPseq k-means clustering analysis across developmental stages, related to Figure 7. **(A)** Heatmaps showing k-means clustering results based on normalized projection strengths to five target regions at three developmental stages: P14→P16, P22→P24, and P60+→P62+. **(B)** Cluster evaluation using four methods (Elbow, Silhouette, Gap Statistic, and Bayesian Information Criterion (BIC)) to determine the optimal number of clusters for each age group, with the consensus estimate indicated (red dotted lines).

**Figure S6.**
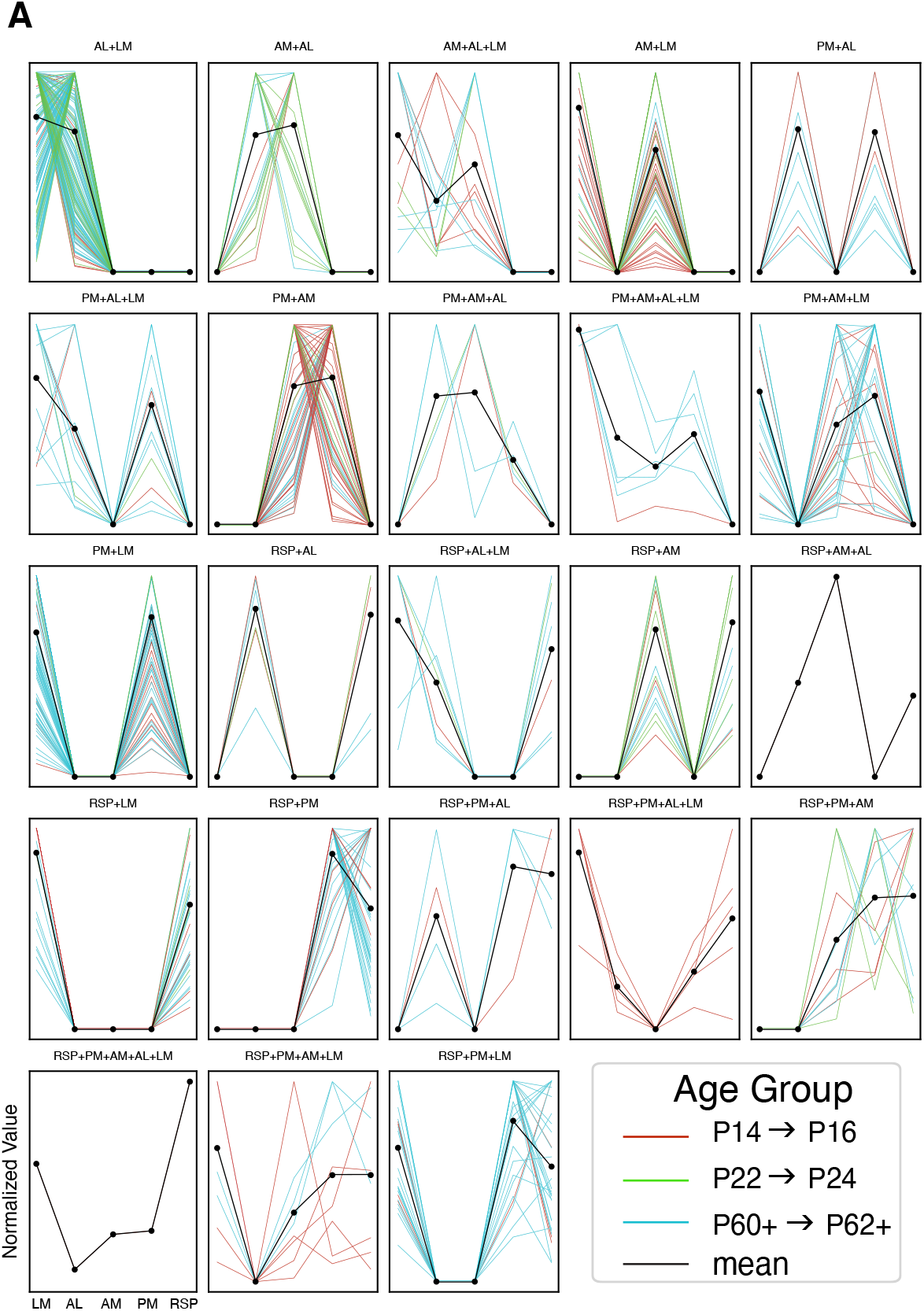
Comprehensive dataset of projection strengths from individual MAPseq-labeled neurons, showing developmental changes across projection motifs and three postnatal stages, related to Figure 7. **(A)** Projection strengths of individual neurons for 23 representative motifs at three developmental time points: P14→P16 (red), P22→P24 (green), and P60+→P62+ (blue). Black points indicate the mean projection strength for each target area.

## SUPPLEMENTAL TABLES

Supplemental tables were provided separately as excel files in a zip file.

**Table S1. Summary of animals and experimental conditions, related to Figures 1-7**.

**Table S2. Summary of raw cell-count data from VSVdG and AAVretro-H2B tracing experiments, related to Figures 2**,**3**,**5**,**6**.

**Table S3. Summary of statistical details for VSVdG and AAVretro-H2B tracing experiments, related to Figures 2**,**3**,**5**,**6**.

**Table S4. Summary of raw cell-count data from axonal pruning versus apoptosis experiments, related to Figure 4**.

**Table S5. Summary of MAPseq motif distribution statistics, related to Figure 7**.

## REFERENCES

1. Felleman, D.J., and Van Essen, D.C. (1991). Distributed hierarchical processing in the primate cerebral cortex. Cereb. Cortex 1, 1–47. 10.1093/cercor/1.1.1-a.

2. Harris, J.A., Mihalas, S., Hirokawa, K.E., Whitesell, J.D., Choi, H., Bernard, A., Bohn, P., Caldejon, S., Casal, L., Cho, A., et al. (2019). Hierarchical organization of cortical and thalamic connectivity. Nature 575, 195–202. 10.1038/s41586-019-1716-z.

3. Zeki, S., and Shipp, S. (1988). The functional logic of cortical connections. Nature 335, 311–317. 10.1038/335311a0.

4. Baker, J.T., Holmes, A.J., Masters, G.A., Yeo, B.T.T., Krienen, F., Buckner, R.L., and Öngür, D. (2014). Disruption of cortical association networks in schizophrenia and psychotic bipolar disorder. JAMA Psychiatry 71, 109–118. 10.1001/jamapsychiatry.2013.3469.

5. Haberl, M.G., Zerbi, V., Veltien, A., Ginger, M., Heerschap, A., and Frick, A. (2015). Structural-functional connectivity deficits of neocortical circuits in the Fmr1 (-/y) mouse model of autism. Sci. Adv. 1, e1500775. 10.1126/sciadv.1500775.

6. Wang, Q., and Burkhalter, A. (2007). Area map of mouse visual cortex. J. Comp. Neurol. 502, 339–357. 10.1002/cne.21286.

7. Kim, E.J., Zhang, Z., Huang, L., Ito-Cole, T., Jacobs, M.W., Juavinett, A.L., Senturk, G., Hu, M., Ku, M., Ecker, J.R., et al. (2020). Extraction of Distinct Neuronal Cell Types from within a Genetically Continuous Population. Neuron 107, 274–282.e6. 10.1016/j.neuron.2020.04.018.

8. Glickfeld, L.L., Andermann, M.L., Bonin, V., and Reid, R.C. (2013). Cortico-cortical projections in mouse visual cortex are functionally target specific. Nat. Neurosci. 16, 219–226. 10.1038/nn.3300.

9. Marshel, J.H., Garrett, M.E., Nauhaus, I., and Callaway, E.M. (2011). Functional specialization of seven mouse visual cortical areas. Neuron 72, 1040–1054. 10.1016/j.neuron.2011.12.004.

10. Han, Y., Kebschull, J.M., Campbell, R.A.A., Cowan, D., Imhof, F., Zador, A.M., and Mrsic-Flogel, T.D. (2018). The logic of single-cell projections from visual cortex. Nature 556, 51–56. 10.1038/nature26159.

11. Young, H., Belbut, B., Baeta, M., and Petreanu, L. (2021). Laminar-specific cortico-cortical loops in mouse visual cortex. Elife 10. 10.7554/eLife.59551.

12. Kast, R.J., and Levitt, P. (2019). Precision in the development of neocortical architecture: From progenitors to cortical networks. Prog. Neurobiol. 175, 77–95. 10.1016/j.pneurobio.2019.01.003.

13. Innocenti, G.M., and Price, D.J. (2005). Exuberance in the development of cortical networks. Nat. Rev. Neurosci. 6, 955–965. 10.1038/nrn1790.

14. Luo, L., and O’Leary, D.D.M. (2005). Axon retraction and degeneration in development and disease. Annu. Rev. Neurosci. 28, 127–156. 10.1146/annurev.neuro.28.061604.135632.

15. Innocenti, G.M., Fiore, L., and Caminiti, R. (1977). Exuberant projection into the corpus callosum from the visual cortex of newborn cats. Neurosci. Lett. 4, 237–242. 10.1016/0304-3940(77)90185-9.

16. Katz, L.C., and Shatz, C.J. (1996). Synaptic activity and the construction of cortical circuits. Science 274, 1133–1138. 10.1126/science.274.5290.1133.

17. Sanes, J.R., and Lichtman, J.W. (1999). Development of the vertebrate neuromuscular junction. Annu. Rev. Neurosci. 22, 389–442. 10.1146/annurev.neuro.22.1.389.

18. Callaway, E.M., and Katz, L.C. (1990). Emergence and refinement of clustered horizontal connections in cat striate cortex. J. Neurosci. 10, 1134–1153. 10.1523/JNEUROSCI.10-04-01134.1990.

19. Klingler, E., Tomasello, U., Prados, J., Kebschull, J.M., Contestabile, A., Galiñanes, G.L., Fièvre, S., Santinha, A., Platt, R., Huber, D., et al. (2021). Temporal controls over interareal cortical projection neuron fate diversity. Nature 599, 453–457. 10.1038/s41586-021-04048-3.

20. Molyneaux, B.J., Arlotta, P., Menezes, J.R.L., and Macklis, J.D. (2007). Neuronal subtype specification in the cerebral cortex. Nat. Rev. Neurosci. 8, 427–437. 10.1038/nrn2151.

21. Barone, P., Dehay, C., Berland, M., and Kennedy, H. (1996). Role of directed growth and target selection in the formation of cortical pathways: prenatal development of the projection of area V2 to area V4 in the monkey. J. Comp. Neurol. 374, 1–20. 10.1002/(SICI)1096-9861(19961007)374:1<1::AID-CNE1>3.0.CO;2-7.

22. Kebschull, J.M., Garcia da Silva, P., Reid, A.P., Peikon, I.D., Albeanu, D.F., and Zador, A.M. (2016). High-throughput mapping of single-neuron projections by sequencing of barcoded RNA. Neuron 91, 975–987. 10.1016/j.neuron.2016.07.036.

23. Han, X., and Bonin, V. (2024). Higher-order cortical and thalamic pathways shape visual processing streams in the mouse cortex. Curr. Biol. 10.1016/j.cub.2024.10.048.

24. Cheng, S., Butrus, S., Tan, L., Xu, R., Sagireddy, S., Trachtenberg, J.T., Shekhar, K., and Zipursky, S.L. (2022). Vision-dependent specification of cell types and function in the developing cortex. Cell 185, 311–327.e24. 10.1016/j.cell.2021.12.022.

25. Soronow, A.L.R., Jacobs, M.W., Dickson, R.G., and Kim, E.J. (2025). Bell Jar: A semiautomated registration and cell counting tool for mouse neurohistology analysis. eNeuro 12. 10.1523/ENEURO.0036-23.2025.

26. Kronman, F.N., Liwang, J.K., Betty, R., Vanselow, D.J., Wu, Y.-T., Tustison, N.J., Bhandiwad, A., Manjila, S.B., Minteer, J.A., Shin, D., et al. (2024). Developmental mouse brain common coordinate framework. Nat. Commun. 15, 9072. 10.1038/s41467-024-53254-w.

27. Wang, Q., Ding, S.-L., Li, Y., Royall, J., Feng, D., Lesnar, P., Graddis, N., Naeemi, M., Facer, B., Ho, A., et al. (2020). The Allen Mouse Brain Common Coordinate Framework: A 3D reference atlas. Cell 181, 936–953.e20. 10.1016/j.cell.2020.04.007.

28. Li, Z., Li, P., Pan, H., Adams, A.M.J., Huang, Z., Liu, H., Li, J., Su, Y., Zhong, B., Bray, L., et al. (2025). D-LMBmapX: Generalized deep-learning pipeline for automated wholebrain neural circuitry profiling across any developmental stages. bioRxiv. 10.1101/2025.02.25.639766.

29. Liwang, J.K., Kronman, F.A., Minteer, J.A., Wu, Y.-T., Vanselow, D.J., Ben-Simon, Y., Taormina, M., Parmaksiz, D., Way, S.W., Zeng, H., et al. (2023). epDevAtlas: Mapping GABAergic cells and microglia in postnatal mouse brains. bioRxivorg. 10.1101/2023.11.24.568585.

30. Gămănuţ, R., Kennedy, H., Toroczkai, Z., Ercsey-Ravasz, M., Van Essen, D.C., Knoblauch, K., and Burkhalter, A. (2018). The mouse cortical connectome, characterized by an ultradense cortical graph, maintains specificity by distinct connectivity profiles. Neuron 97, 698–715.e10. 10.1016/j.neuron.2017.12.037.

31. Wang, Q., Gao, E., and Burkhalter, A. (2011). Gateways of ventral and dorsal streams in mouse visual cortex. J. Neurosci. 31, 1905–1918. 10.1523/JNEUROSCI.3488-10.2011.

32. Cai, L., Argunşah, A.Ö., Damilou, A., and Karayannis, T. (2024). A nasal chemosensationdependent critical window for somatosensory development. Science 384, 652–660. 10.1126/science.adn5611.

33. Markov, N.T., Vezoli, J., Chameau, P., Falchier, A., Quilodran, R., Huissoud, C., Lamy, C., Misery, P., Giroud, P., Ullman, S., et al. (2014). Anatomy of hierarchy: feedforward and feedback pathways in macaque visual cortex. J. Comp. Neurol. 522, 225–259. 10.1002/cne.23458.

34. Watakabe, A., Skibbe, H., Nakae, K., Abe, H., Ichinohe, N., Rachmadi, M.F., Wang, J., Takaji, M., Mizukami, H., Woodward, A., et al. (2023). Local and long-distance organization of prefrontal cortex circuits in the marmoset brain. Neuron 0. 10.1016/j.neuron.2023.04.028.

35. Siu, C., Balsor, J., Merlin, S., Federer, F., and Angelucci, A. (2021). A direct interareal feedback-to-feedforward circuit in primate visual cortex. Nat. Commun. 12, 4911. 10.1038/s41467-021-24928-6.

36. Hand, R.A., Khalid, S., Tam, E., and Kolodkin, A.L. (2015). Axon Dynamics during Neocortical Laminar Innervation. Cell Rep. 12, 172–182. 10.1016/j.celrep.2015.06.026.

37. Garrett, M.E., Nauhaus, I., Marshel, J.H., and Callaway, E.M. (2014). Topography and areal organization of mouse visual cortex. J. Neurosci. 34, 12587–12600. 10.1523/JNEUROSCI.1124-14.2014.

38. Watakabe, A., and Hirokawa, J. (2018). Cortical networks of the mouse brain elaborate within the gray matter. Brain Struct. Funct. 223, 3633–3652. 10.1007/s00429-018-1710-5.

39. Yasuda, M., Nagappan-Chettiar, S., Johnson-Venkatesh, E.M., and Umemori, H. (2021). An activity-dependent determinant of synapse elimination in the mammalian brain. Neuron 109, 1333–1349.e6. 10.1016/j.neuron.2021.03.006.

40. Smith, I.T., Townsend, L.B., Huh, R., Zhu, H., and Smith, S.L. (2017). Stream-dependent development of higher visual cortical areas. Nat. Neurosci. 20, 200–208. 10.1038/nn.4469.

41. Murakami, T., Matsui, T., and Ohki, K. (2017). Functional segregation and development of mouse higher visual areas. J. Neurosci. 37, 9424–9437. 10.1523/JNEUROSCI.0731-17.2017.

42. Murakami, T., Matsui, T., Uemura, M., and Ohki, K. (2022). Modular strategy for development of the hierarchical visual network in mice. Nature 608, 578–585. 10.1038/s41586-022-05045-w.

43. Chen, X., Fischer, S., Rue, M.C.P., Zhang, A., Mukherjee, D., Kanold, P.O., Gillis, J., and Zador, A.M. (2024). Whole-cortex in situ sequencing reveals input-dependent area identity. Nature. 10.1038/s41586-024-07221-6.

44. Berezovskii, V.K., Nassi, J.J., and Born, R.T. (2011). Segregation of feedforward and feedback projections in mouse visual cortex. J. Comp. Neurol. 519, 3672–3683. 10.1002/cne.22675.

45. Liu, S., Gao, L., Chen, J., and Yan, J. (2024). Single-neuron analysis of axon arbors reveals distinct presynaptic organizations between feedforward and feedback projections. Cell Rep. 43, 113590. 10.1016/j.celrep.2023.113590.

46. Robinson, S., Poorman, C.E., Marder, T.J., and Bucci, D.J. (2012). Identification of functional circuitry between retrosplenial and postrhinal cortices during fear conditioning. J. Neurosci. 32, 12076–12086. 10.1523/JNEUROSCI.2814-12.2012.

47. Ward, J.H., Jr (1963). Hierarchical grouping to optimize an objective function. J. Am. Stat. Assoc. 58, 236–244. 10.1080/01621459.1963.10500845.

48. Rousseeuw, P.J. (1987). Silhouettes: A graphical aid to the interpretation and validation of cluster analysis. J. Comput. Appl. Math. 20, 53–65. 10.1016/0377-0427(87)90125-7.

49. Calinski, T., and Harabasz, J. (1974). A dendrite method for cluster analysis. Commun. Stat. Theory Methods 3, 1–27. 10.1080/03610927408827101.

50. Davies, D.L., and Bouldin, D.W. (1979). A Cluster Separation Measure. IEEE Trans. Pattern Anal. Mach. Intell. PAMI-1, 224–227. 10.1109/tpami.1979.4766909.

51. 1 | 1967 Some methods for classification and analysis of multivariate observations Chapter Author(s) (1967). Jerzy Neyman Berkeley Symp. on Math. Statist. and Prob 5, 281–297.

52. Oh, S.W., Harris, J.A., Ng, L., Winslow, B., Cain, N., Mihalas, S., Wang, Q., Lau, C., Kuan, L., Henry, A.M., et al. (2014). A mesoscale connectome of the mouse brain. Nature 508, 207–214. 10.1038/nature13186.

53. Bridle, J. (1989). Training stochastic model recognition algorithms as networks can lead to maximum mutual information estimation of parameters. Neural Inf Process Syst, 211–217.

54. Majtey, A.P., Lamberti, P.W., and Prato, D.P. (2005). Jensen-Shannon divergence as a measure of distinguishability between mixed quantum states. Phys. Rev. A 72, 052310. 10.1103/PhysRevA.72.052310.

